# Exploiting De Novo Serine Synthesis as a Metabolic Vulnerability to Overcome Sunitinib Resistance in Advanced Renal Cell Carcinoma

**DOI:** 10.1101/2024.03.22.586287

**Authors:** Manon Teisseire, Umakant Sahu, Julien Parola, Meng-Chen Tsai, Valérie Vial, Jérôme Durivault, Renaud Grépin, Yann Cormerais, Gilles Pagès, Issam Ben-Sahra, Sandy Giuliano

## Abstract

Sunitinib, an oral tyrosine kinase inhibitor used in advanced renal cell carcinoma (RCC), exhibits significant efficacy but faces resistance in 30% of patients. Yet, the molecular mechanisms underlying this therapy resistance remain elusive. Here, we show that sunitinib induces a metabolic shift leading to increased serine synthesis in RCC cells. The activation of the GCN2-ATF4 stress response pathway is identified as the mechanistic link between sunitinib treatment and elevated serine production. Inhibiting key enzymes in the serine synthesis pathway, such as PHGDH and PSAT1, enhances the sensitivity of resistant cells to sunitinib. The study underscores the role of serine biosynthesis in nucleotide synthesis, influencing cell proliferation, migration, and invasion. Beyond RCC, similar activation of serine synthesis occurs in other cancer types, suggesting a shared adaptive response to sunitinib therapy. This research identifies serine synthesis as a potential target to overcome sunitinib resistance, offering insights into therapeutic strategies applicable across diverse cancer contexts.

**Graphical abstract:** 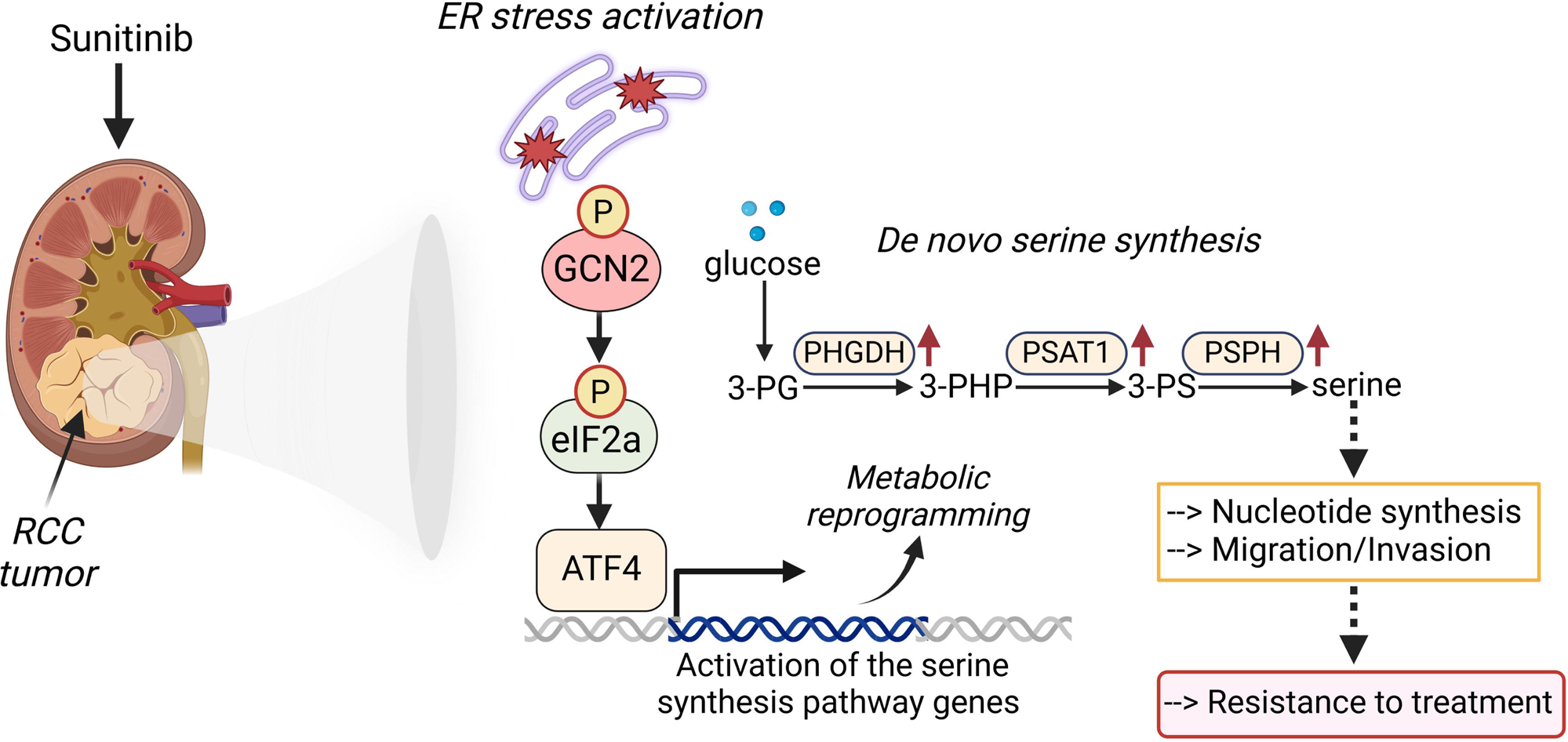

**Highlights:**

- Sunitinib induces an increase in endogenous serine production in metastatic ccRCC.
- The heightened serine biosynthesis promoted by sunitinib facilitates nucleotide synthesis, thereby sustaining tumor cell proliferation.
- Sunitinib-induced enhancement of serine biosynthesis enables cell migration and invasion.
- The stimulation in serine synthesis is also observed in other cancer models treated with sunitinib.

## Introduction

Clear cell renal cell carcinoma (ccRCC), which accounts for 80% of kidney cancer cases, is primarily linked to Von Hippel–Lindau (VHL) and polybromo-1 (PBRM-1) gene abnormalities (^1,2,3^). VHL inactivation stabilizes the hypoxia-inducible factor 1α and 2α (HIF1α/ HIF2α), promoting angiogenesis, migration, invasion, and metabolic reprogramming. Factors such as smoking, dialysis dependence, prolonged painkiller use, high blood pressure, obesity, and diabetes often contribute to VHL and PBRM1 mutations. The deletion of VHL or inactivation of PBRM1 has been shown to promote glycolysis, pro-growth signaling, and cholesterol biosynthesis in tumor cells (PMID: 27100670; PMID: 35672925, PMID: 25890500). Notably, cancer cells exhibit significant glucose consumption, facilitating ATP production and supporting the synthesis of metabolic intermediates such as dihydroxyacetone-phosphate, serine, glycine, and ribose 5-phoshate, which are needed to produce macromolecules including lipids and nucleotides. Increased de novo nucleotide synthesis in cancer cells is crucial for DNA replication and RNA production, sustaining protein synthesis across various stages cell cycle stages (8) (PMID: 3696730).

In the late 2000s, metastatic ccRCC exhibited a grim prognosis, characterized by a median survival of approximately 13 months and a 5-year survival rate below 10%. This challenging outlook primarily stemmed from the limited efficacy of traditional chemotherapy and radiotherapy. The therapeutic landscape for ccRCC has since witnessed improvement with the introduction of anti-angiogenic drugs and immunotherapies, resulting in enhanced survival rates. Despite these advancements, metastatic ccRCC remains an incurable condition and often leads to fatalities (^4^). In numerous countries, the standard of care for metastatic ccRCC relies on well-established tyrosine kinase inhibitors, notably sunitinib and pazopanib. These agents, considered conventional or “ancient” in their inception, remain integral components of the therapeutic armamentarium due to their proven efficacy.

Sunitinib targets the vascular endothelial growth factor receptor (VEGFR) and other tyrosine kinase receptors. However, resistance inevitably develops, with proposed mechanisms including the restoration of angiogenesis (^5,6,7^) and altered drug bioavailability (^8^). Despite these insights, the precise molecular causes of sunitinib resistance in ccRCC remain incompletely understood. It has been demonstrated that ccRCC exhibits dynamic metabolism, characterized by lipid and glycogen accumulation due to HIF1-α-driven aerobic glycolysis (Metabolic reprogramming in clear cell renal cell carcinoma, Hiromi I Wettersten 1, Omran Abu Aboud 2, Primo N Lara Jr 3, Robert H Weiss 2, 2017). Yet, the alteration of other metabolic pathways in ccRCC and whether a specific pathway contributes to sunitinib resistance remain to be demonstrated. Another intriguing effect of sunitinib on ccRCC is its inhibitory impact on autophagy, a key catabolic process that maintains macromolecular equilibrium (^8^). An outstanding question arises: how do cells survive and sustain growth and proliferation without complete autophagy? We hypothesize that ccRCC cells treated with sunitinib adapt by maintaining anabolic metabolism through compensatory pathways, thereby developing resistance to these compounds.

The serine-glycine one-carbon cycle, which arises from glycolysis, generates carbon units and precursors that are critical for nucleotide synthesis (^31,32^), transsulfuration, glutathione metabolism, anabolic and redox processes, and methylation reactions that shape the epigenetic landscape. One-carbon metabolism, considered vital in cancer, produces biosynthetic substrates and cofactors necessary for cancer cell proliferation (^10,11^). Previous studies highlight the critical role of the serine biosynthesis pathway in various cancers (^11–16^) (PMID: 35577785), with enzymes like phosphoglycerate dehydrogenase (PHGDH) being amplified or overexpressed (^16,17,18,19^).

Through the integration of RNA sequencing and metabolite profiling data, we identified a distinct metabolic adaptive response to sunitinib treatment, observed in both sunitinib-sensitive and -resistant ccRCC cells. Our results show a significant accumulation of serine and glycine in ccRCC cells undergoing sunitinib treatment. This accumulation stems from an increased endogenous synthesis of serine and glycine, enabling ccRCC cells to partially evade the inhibitory effects of sunitinib and thereby facilitating tumor progression and resistance. Beyond ccRCC, we demonstrate that sunitinib induced an increase in serine synthesis in diverse cancer models, including triple-negative breast cancer, lung cancer, medulloblastoma, and head and neck cancer. Mechanistically, the effects of sunitinib are mediated by the activation of the GCN2-ATF4 signaling axis within the integrated stress response (ISR) pathway. This activation subsequently upregulates serine synthesis enzymes, propelling an increase in serine and glycine production. The induction of serine synthesis serves as a compensatory mechanism, enabling cells to overcome autophagy defects and support nucleotide synthesis for sustained proliferation and tumor progression. Importantly, direct blockade of serine biosynthesis not only impedes tumor growth but also restricts cell migration and invasion. Our findings suggest that the induction of serine synthesis is a metabolic hallmark of sunitinib resistance, highlighting its potential as a therapeutic target in advanced ccRCC.

## RESULTS

### Sunitinib leads to an upregulation of the serine biosynthesis pathway in ccRCC cells

To investigate the transcriptional alterations induced by sunitinib in ccRCC, we subjected the renal cancer cell line 786-O to a 48-hour treatment with sunitinib, followed by RNA sequencing analysis. Notably, the analysis revealed 283 downregulated and 478 upregulated transcripts in sunitinib-treated cells (Figure S1A).

To elucidate the functions of differentially expressed genes in sunitinib-treated cells, we employed gene set enrichment analysis (GSEA) and conducted functional analyses using gene ontology (GO) terms and the Kyoto Encyclopedia of Genes and Genomes (KEGG) pathway computational tool (Figure S1B). Sunitinib treatment led to a significant enrichment of multiple KEGG pathways, including glycine, serine, and threonine metabolism, PI3K-Akt signaling pathway, p53 signaling pathway, lysosome, and cell cycle (Figure 1A, and Figure S1C). The glycine, serine and threonine pathway is one of the most highly activated metabolic pathways shared by two RCC cell lines, the 786-O and A498 cell lines (Figure S1D).

**Figure 1:**
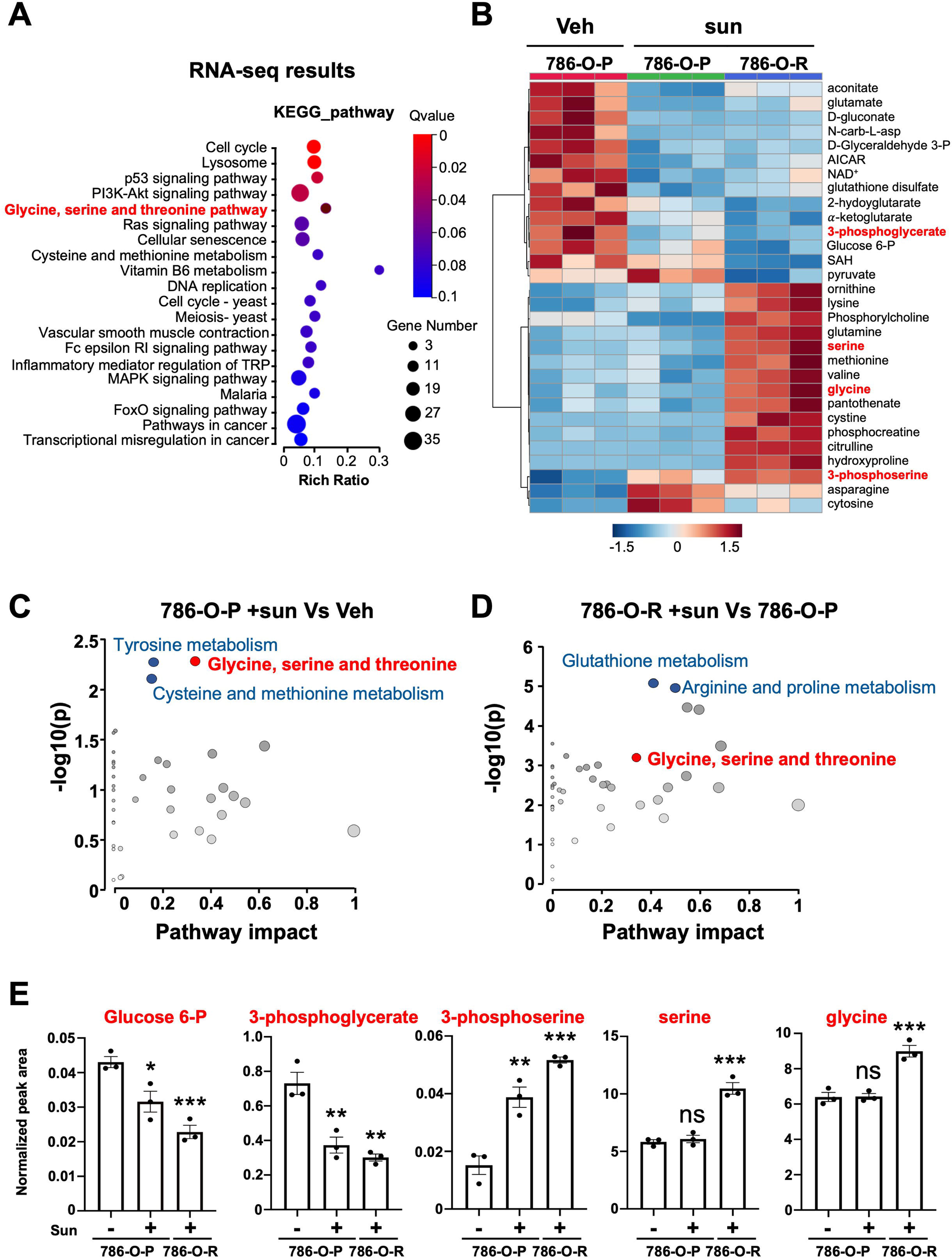
Sunitinib-treated mccRCC shows metabolic changes and increases in glycine, serine, and threonine metabolism. (A) Functional enrichment results based on KEGG were presented as dot-bubble diagrams (pathways with ascending significance from top to bottom) in sunitinib-treated 786-O cells compared to 786-O control cells. Bubble area is proportional to the effect of the individual metabolic pathways. The most strongly altered metabolic pathways are characterized by both a high log(p) value and a high impact value (upper right region. Glycine, serine, and threonine metabolism are shown in red). (B) Steady-state metabolite profiles of 786-O cells treated with DMSO (vehicle, veh) or sunitinib (sun, 2.5μM, 48h). Intracellular metabolites from three independent samples per condition were profiled by LCMS/MS, and the significantly elevated metabolites are shown as line-normalized heatmaps ordered by p-value. Heatmap visualization of the 30 major metabolites differentially expressed in 786-O parental cells (786-O-P) undercontrol (vehicle, veh), sunitinib-treated (sun, 2.5μM, 16h) and sunitinib-resistant (786-O-R) treated with sunitinib (sun, 2.5μM, 16h), with each condition in triplicate. (C) Bubble plots depict altered metabolic pathways in 786-O parental cells treated with sunitinib (786-O P +sun) compared to DMSO (veh). (D) Bubble plots illustrate the differences in altered metabolic pathways between sunitinib-resistant cells (786-O R) and parental cells (786-O-P). (E) Normalized peak areas of various metabolites serving as intermediates in the serine biosynthesis pathway are presented to provide insights into the impact of sunitinib treatment and resistance mechanisms. ns, non-significant; * p < 0.05; ** p < 0.01; *** p < 0.001 (two-way ANOVA).

To understand whether changes in metabolic gene expression corresponded to alterations at the metabolite level, we conducted targeted metabolomic analysis on parental 786-O cells, sunitinib-sensitive 786-O cells, and sunitinib-resistant 786-O cells, treated with vehicle or sunitinib (Figure S2A-B). The analysis revealed differential production of various metabolites, including intermediates in de novo serine/glycine synthesis (Figure 1D-E, Figure S2C).

Among the altered metabolic pathways, glycine, serine, and threonine metabolism stood out as the most upregulated network in sunitinib-sensitive (Figure 1C, Figure S2D; impact factor≥0.1) and resistant cells (Figure 1D, Figure S2E; impact factor≥0.1).

Accumulation of serine/glycine biosynthesis intermediates, such as 3-phosphoserine, serine, and glycine, was observed in cells treated with sunitinib for 48 hours. This accumulation was more pronounced in sunitinib-resistant cells compared to the parent 786-O cells treated with a vehicle (Figure 1E).

Proliferating cells synthesize serine from the glycolytic intermediate 3-phosphoglycerate (3PG) through a three-step enzymatic process, involving phosphoglycerate dehydrogenase (PHGDH), phosphoserine aminotransferase 1 (PSAT1), and phosphoserine phosphatase (PSPH) (Figure 2A) (PMID: 24657017, PMID: 23822983).

**Figure 2:**
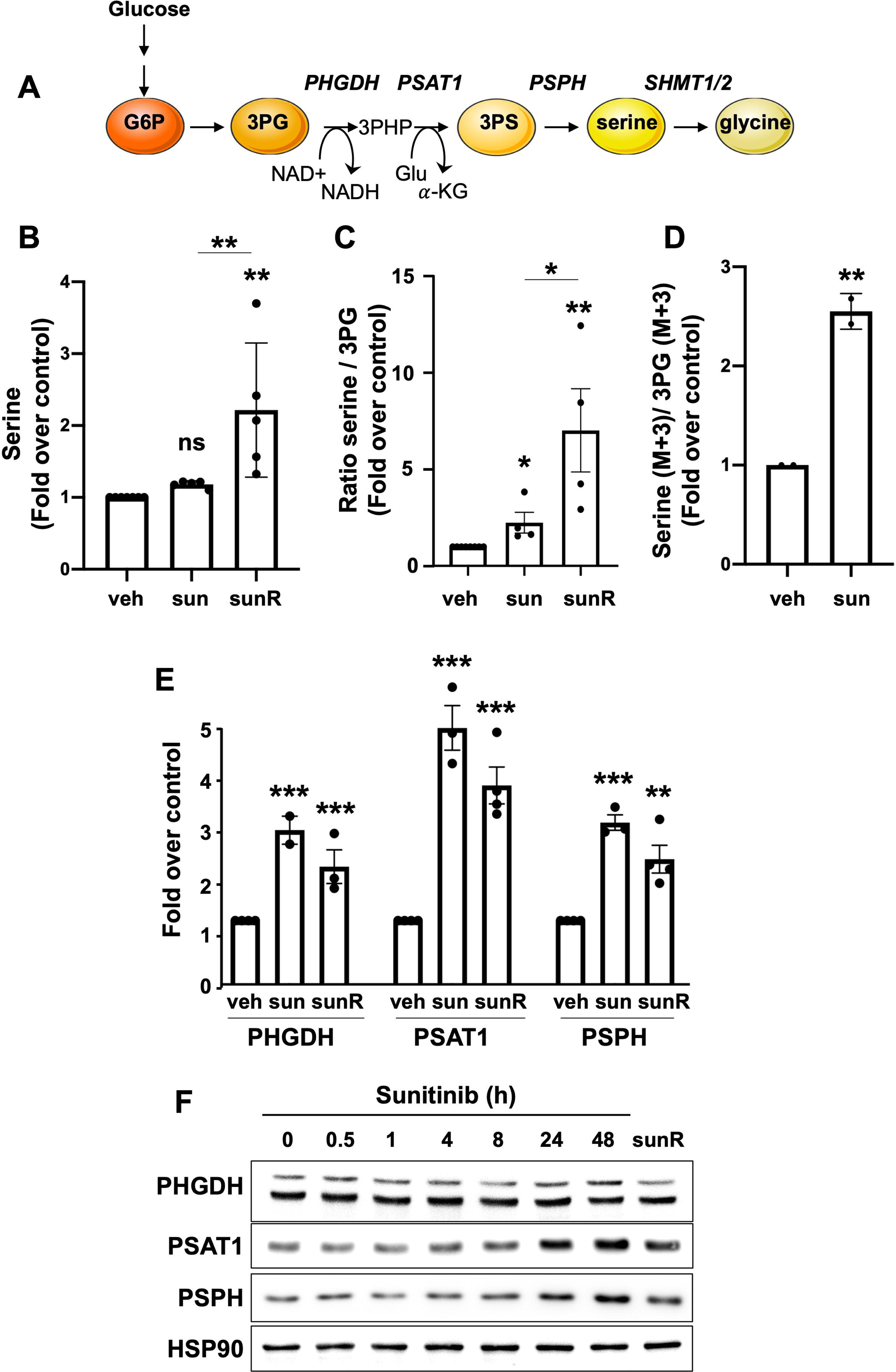
Treatment with sunitinib leads to an increase in the serine biosynthesis pathway in vitro. (A) Schematic representation of the serine biosynthesis pathway and the three enzymes directly involved and associated with this pathway. (B) Normalized peak areas of serine production from five independent steady-state experiments measured by targeted LCMS/MS of RCC 786-O cells as control (vehicle, veh), sunitinib-treated cells (sun, 2.5μM, 48h) and sunitinib-resistant cells (R). (C) Ratio of serine / 3-phosphoglycerate production of four independent steady-state experiments measured by targeted LCMS/MS of RCC 786-O cells as control (veh), sunitinib-treated cells (sun, 2.5μM, 48h) and sunitinib-resistant cells measured (sunR). (D) Ratio of serine / 3-phosphoglycerate production of two independent flux experiments measured by LCMS/MS of RCC 786-O cells control, sunitinib treated cells (2.5*μ*;M, 48h) after incubation in ^13^C^6^ glucose-containing culture medium for 16 hours. (E) mRNA levels of PHGDH, PSAT1 and PSPH from RCC 786-O treated with sunitinib (sun, 2.5μM, 48h) or cells resistant to this treatment (R). (F) Immunoblots of RCC 786-O cells treated with sunitinib at different times or cells resistant to this treatment. ns, non-significant; * p < 0.05; ** p < 0.01; *** p < 0.001 (two-way ANOVA).

A significant increase in serine levels and the serine/3PG ratio was observed in sunitinib-resistant cells (Figure 2B-C). Stable isotope tracing using [^13^C_6_]-glucose confirmed an increase in the rate of serine synthesis in sunitinib-treated 786-O cells compared to that of vehicle-treated 786-O cells (Figure 2D).

Significantly, *PHGDH*, *PSPH*, and *PSAT1* mRNA expression increased following both short-term sunitinib treatment and chronic exposure, relative to control counterparts (Figure 2E). The increase of *PSAT1* and *PSPH* mRNA expression under sunitinib treatment and in resistant cells was confirmed at the protein levels, but PHGDH protein levels remained unchanged (Figure 2F).

Cytosolic *SHMT1* and mitochondrial *SHMT2*, responsible for converting serine to glycine and generating an activated one-carbon unit, exhibited increased mRNA levels, while protein levels remained unaltered in 786-O cells (Figures S3A and S3B) and RCC10 cells (Figure S3C-E).

Overall, these gene expression and metabolic profiles reveal an upregulation of the serine/glycine synthesis pathway in renal cancer cells in response to sunitinib treatment, underscoring the interplay between transcriptional regulation and metabolic adaptation in the context of sunitinib treatment and resistance.

### The induction of the integrated stress response (ISR) pathway is required for sunitinib-dependent de novo serine synthesis

Amino acid and glucose deprivation, along with the generation of reactive oxygen species (ROS), which are common stress factors in solid tumors, activate the eukaryotic initiation factor 2α (eIF2α)-kinase GCN2 (^20,21,22^). Once activated, GCN2 phosphorylates eIF2α, leading to the translational upregulation of activating transcription factor 4 (ATF4) (^23,24^). ATF4 is the main transcription factor known to directly bind the promoter of serine synthesis genes, subsequently promoting their expression (^25,26^). Upon conditions such as amino acid starvation, eIF2α phosphorylation reduces global protein synthesis but selectively enables the translation of certain genes including ATF4, facilitating metabolic adaptation and cell survival (^23,24,27^).

The ISR pathway was activated within 30 minutes of sunitinib treatment in 786-O cells, evidenced by increased phosphorylation of GCN2, eIF2α, and induction of ATF4 (Figure 3A-B). As a control, we induced ISR with tunicamycin, yielding similar effects on the eIF2α-ATF4 axis and increasing PSAT1 and PSPH protein levels (Figure 3B). The regulation of GCN2-eIF2α and the increase in serine and glycine levels upon both sunitinib and tunicamycin conditions in sensitive or resistant cells (Figure 3C), illustrate the canonical induction of ISR under these conditions. To determine whether GCN2 and ATF4 are required for increased serine synthesis enzymes under sunitinib treatment, we suppressed *GCN2* (Figure 3D) or *ATF4* (Figure 3E) expression using small interfering RNA (siRNA). Knockdown of *GCN2* or *ATF4* prevented the sunitinib-dependent increase in PSAT1 and PSPH protein levels. In contrast, no changes were observed on SHMT1 and SHMT2 protein levels upon sunitinib treatment and *ATF4* knockdown (Figure S4A). In our model, we propose that the GCN2-eIF2α pathway is required to mediate the sunitinib-dependent induction of ATF4 expression. ATF4 translation is typically induced by the phosphorylation of eIF2α, a process reliant on upstream open reading frames (uORFs) in the 5’ UTR of *ATF4* transcript (^37^). Through cycloheximide protein stability assay, we showed that sunitinib treatment, similarly to tunicamycin, increased ATF4 protein stability in 786-O cells (Figure S4B).

**Figure 3:**
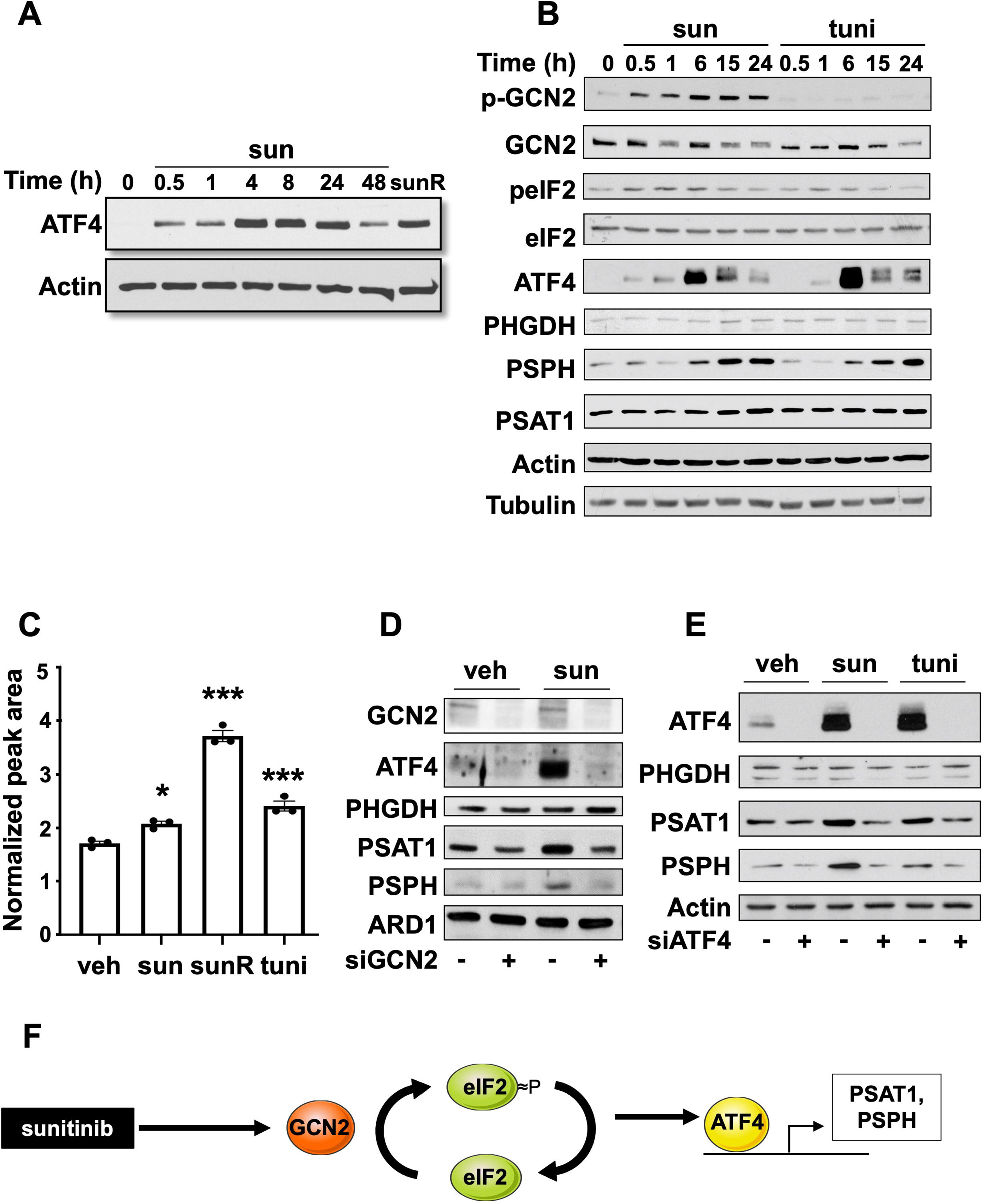
The GCN2/eIF2/ATF4 RE stress response pathway is required for activation of the serine biosynthesis pathway. (A) Protein expression profiles by immunoblots of ATF4 in 786-O Parental ccRCC cells treated with either DMSO (veh) or a time course of sunitinib (sun, 2.5μM) or in 786-O resitant cells (sunR). (B) Protein expression profiles by immunoblots in 786-O ccRCC cells treated with DMSO (veh) or with a time course of sunitinib (sun, 2.5μM) or a time course of tunicamycin (tuni, 1μg/ml). (C) Normalized peak areas of serine from steady-state metabolite analysis of 786-O treated with DMSO (veh), sunitinib (sun, 2.5μM), tunicamycin (tuni, 1μg/ml), or sunitinib-resistant cells (sunR). (D) Protein expression profiles by immunoblots in 786-O RCC cells treated with DMSO (veh) or sunitinib (sun, 2.5μM) and transfected with siRNAs targeting GCN2 (siGCN2) or nontargeting controls for 48 hours. (E) Protein expression profiles by immunoblots of 786-O RCC cells treated with DMSO (veh), sunitinib (sun, 2.5μM) or tunicamycin (tuni, 1μg/ml) and transfected with siRNAs targeting ATF4 (siATF4) or non-targeting controls for 48 hours. (F) Schematic diagram illustrating the mechanism induced by sunitinib treatment. The increase in serine synthesis upon sunitinib treatment requires the activation of the integrative stress response pathway GCN2-ATF4, governing the expression of serine synthesis enzymes. * p < 0.05; *** p < 0.001 (two-way ANOVA).

While mTORC1 signaling has been shown to control ATF4 protein levels independently of the canonical ISR pathway (PMID: 26912861)(^38^), rapamycin treatment did not prevent the induction of ATF4 expression in response to sunitinib. Similarly, inhibition of PKC (GF109203X, GFX) or ERK (Uo126) did not avert the induction of ATF4 upon sunitinib treatment (Figure S4C). Therefore, mTORC1, PKC, and ERK do not contribute to the sunitinib-dependent regulation of ATF4, suggesting that the regulation of ATF4 upon sunitinib treatment is independent of the molecular mechanisms involved in growth-factor-dependent control of ATF4 expression.

Consistent results were observed in RCC10 cells under sunitinib treatment, confirming the activation of the GCN2-eIF2α-ATF4 pathway (Figure S4D). ATF4, PSAT1, and PSPH protein levels were also increased in RCC10 cells treated with tunicamycin (Figure S4D). The role of ATF4 in regulating PSAT1 and PSPH, but not PHGDH, was confirmed using siRNA in RCC10 cells (Figure S4E). In summary, the increased serine synthesis following sunitinib treatment necessitates the activation of the integrated stress response pathway GCN2-eIF2α-ATF4, which orchestrates the expression of key serine synthesis enzymes.

### The induction of de novo serine pathway upon sunitinib supports nucleotide synthesis, cell survival, migration, and invasion

To understand if the synthesis of serine serves as a metabolic mechanism enabling RCC cells to resist sunitinib treatment, we employed NCT-503 (^12^), an inhibitor of PHGDH, the rate-limiting enzyme of de novo serine synthesis. The combination of sunitinib and NCT-503 significantly reduced the proliferation of both sunitinib-sensitive and resistant RCC cells, surpassing the efficacy of individual treatments (Figure 4A). To further validate these observations, we utilized a genetic approach, employing siRNA targeting either *PHGDH*, *PSAT1* or *PSPH* within the serine synthesis pathway in the presence or absence of sunitinib. Knockdown of *PSPH* in combination with sunitinib exhibited the most striking decrease in RCC cell number compared to the knockdown of other serine synthesis genes (Figure 4B).

**Figure 4:**
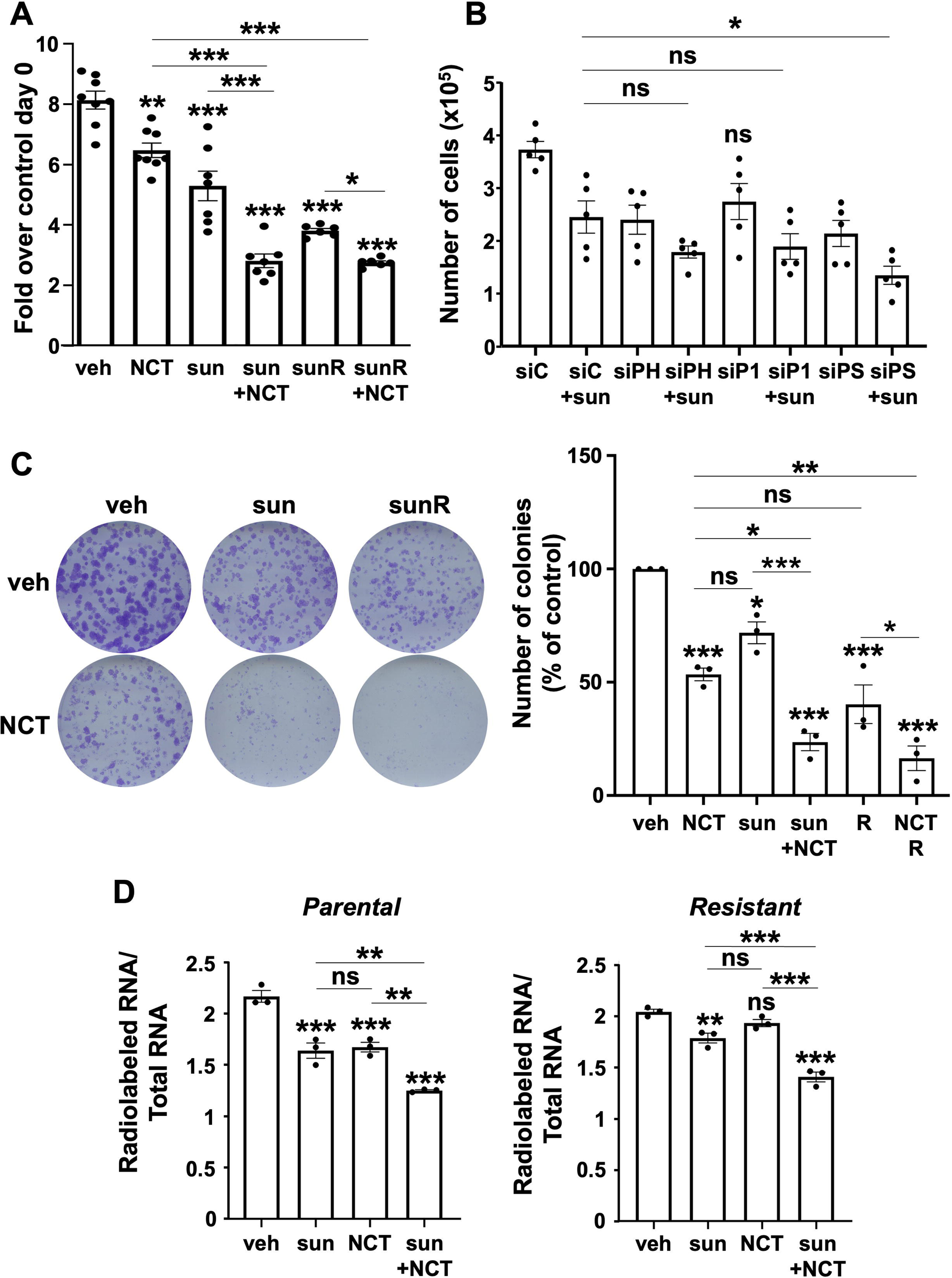
The de novo serine metabolic pathway is essential for proliferation and nucleotide synthesis under sunitinib treatment. (A) CellTiter-Glo measurements performed 96 hours after treating 786-O cells with DMSO (veh), sunitinib (sun, 2.5μM), NCT-503 (NCT, 30μM), or a combination of both treatments. (B) Cell number measurements conducted 96 hours post-treatment of 786-O cells with DMSO (veh) or sunitinib (sun, 2.5μM) and transfected with siRNAs targeting PHGDH (siPH), PSAT1 (siP1), or PSPH (siPS), alongside non-targeted controls (siC) for 96 hours before cell counting. (C) Clonogenic assays performed in 786-O parental cells or sunitinib-resistant cells (sunR) treated with DMSO (veh), sunitinib (sun, 2.5μM), NCT-503 (NCT, 30μM), or a combination of both treatments. Colonies were cultured for 10 days after treatment, photographed, and the colony area was quantified using ImageJ. Three different experiments were conducted. (D) Relative incorporation of radiolabel (C3H3)-methionine into total RNA of parental 786-O cells treated with sunitinib or in resistant cells with DMSO (veh), sunitinib (sun, 2.5μM), NCT-503 (NCT, 30μM) or a combination of both treatments for 24 hours. ns, non-significant; * p < 0.05; ** p < 0.01; *** p < 0.001 (two-way ANOVA).

Furthermore, a colony formation assay revealed a notable reduction in colony formation when sunitinib-sensitive or resistant cells were co-treated with sunitinib and NCT-503, as opposed to single treatments (Figure 4C). These effects were confirmed in a second RCC cell line, RCC10 cells, supporting the critical role of serine synthesis in cell proliferation, particularly under the influence of sunitinib treatment and resistance (Figure S5A). These results suggest potential therapeutic strategies involving the inhibition of this pathway to enhance the efficacy of ccRCC treatment.

The increased serine synthesis induced by sunitinib also led to increased glycine levels in both sunitinib-sensitive and resistant RCC cells (Figure 1E). Considering the role of serine and glycine as substrates for nucleotide synthesis, we hypothesize that RCC cells resist sunitinib treatment by sustaining de novo nucleotide synthesis through serine/glycine synthesis stimulation. To investigate the impact of sunitinib on the metabolic flux of newly synthesized nucleotides into nucleic acids, we measured the incorporation of carbon from U-[^14^C]-glucose into RNA. While sunitinib alone reduced the incorporation of ^14^C from glucose into RNA, the combination of sunitinib and NCT-503 resulted in a further decrease in ^14^C incorporation into RNA in both sensitive and resistant cells compared to NCT-503 or sunitinib treatment alone (Figure 4D). These findings strongly suggest that nucleotide synthesis is, in part, critically dependent on the sunitinib-induced de novo serine synthesis in RCC cells.

While numerous studies associate serine biosynthesis with tumor growth, limited attention has been given to its role in migration, invasion, and metastasis, with existing studies mainly focused on breast cancer (^28,29^). To assess the effects of serine synthesis on ccRCC migration, invasion, or metastasis, we employed siRNA to reduce PSAT1 expression or utilized a PHGDH inhibitor (NCT-503) to inhibit PHGDH activity in ccRCC cells treated with either vehicle or sunitinib.

In our wound-healing assays, we observed that PSAT1 knockdown significantly decreased the motility of 786-O cells, with no discernible difference in sunitinib-treated cells compared to those treated with vehicle. Strikingly, the anti-migratory effects of PSAT1 knockdown were more pronounced in sunitinib-resistant cells than in vehicle-treated cells with reduced PSAT1 expression (Figure 5A). In spheroid invasion assays, the inhibition of PHGDH with NCT-503 or PSAT1 knockdown in 786-O cells significantly impeded the invasiveness of RCC cells. This effect was further enhanced when combined with sunitinib treatment, demonstrating a synergistic impact.

**Figure 5:**
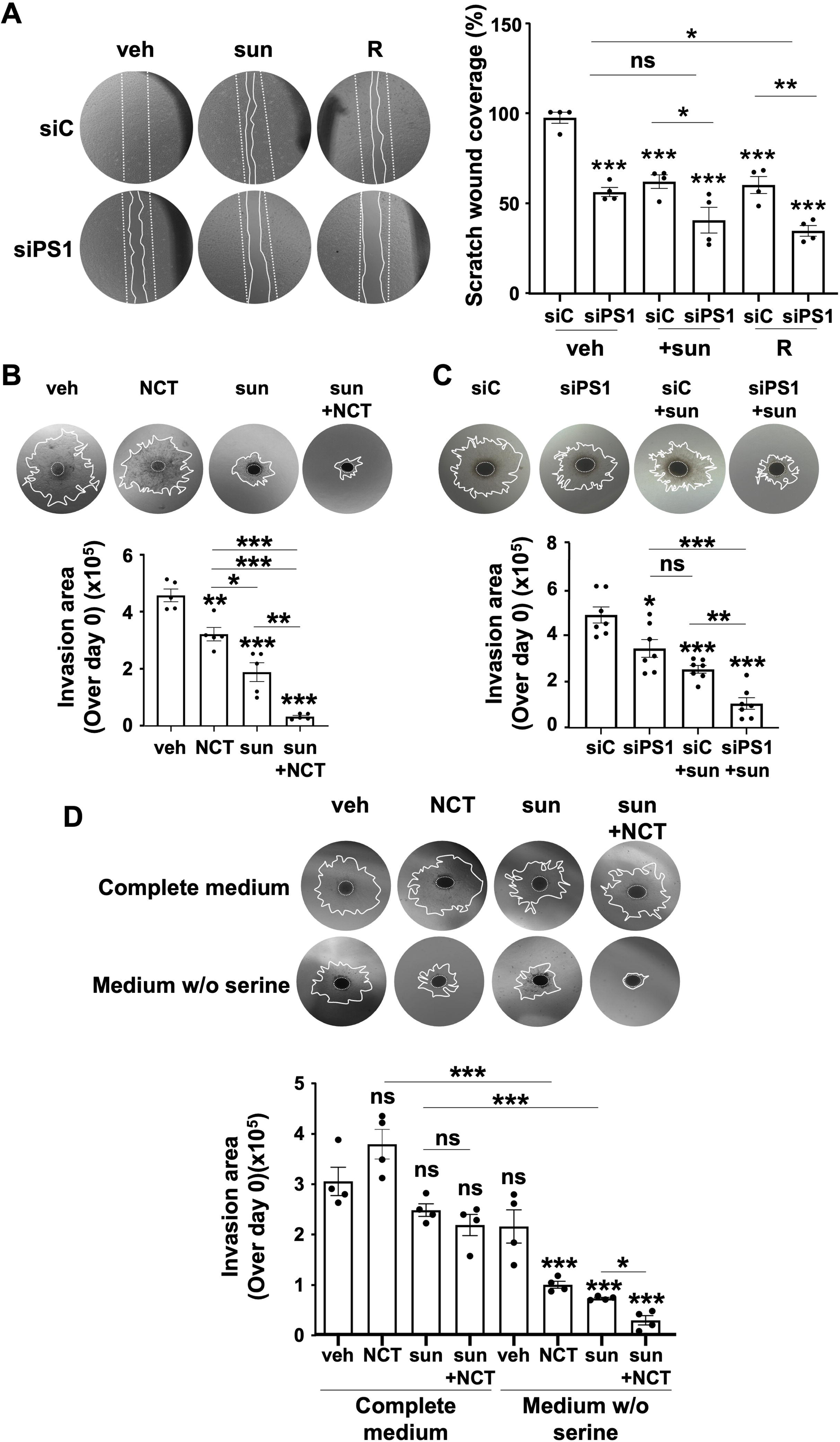
De novo serine signaling pathway is critical for migration and invasion under sunitinib treatment. (A) Cell migration (scratch wound healing experiment). Quantification is shown as percent wound closure ± SEM (n = 4) 16 hours after wounding. (B) Quantification of spheroid invasion area after 4 days of invasion. Spheroids from 786-O parental RCC cells were subsequently treated with DMSO (veh), sunitinib (sun, 2.5μM), NCT-503 (NCT, 30μM), or a combination of both treatments 24 hours before embedding in Matrigel (1 mg/mL). (C) Spheroids were subsequently treated with DMSO or sunitinib (sun, 2.5μM) and transfected with siRNAs targeting PSAT1 (siPSAT1) or nontargeting controls (siC) for 24 hours before embedding in Matrigel (1 mg/mL). (D) spheroids from 786-O resistant RCC cells were subsequently treated with DMSO (veh), sunitinib (sun, 2.5μM), NCT-503 (NCT, 30μM), or a combination of both treatments 24 hours before embedding in Matrigel (1 mg/mL) and cultured in serine-containing or serine-free media. ns, non-significant; * p < 0.05; ** p < 0.01; *** p < 0.001 (two-way ANOVA).

Remarkably, sunitinib-resistant RCC cells cultured in spheroid and exposed to serine-free media exhibited increased sensitivity to PHGDH inhibition (Figure 5D). Both conditions—PHGDH inhibition and serine deprivation in the media— magnified the anti-invasion effects of sunitinib on sunitinib-resistant 786-O cells. In contrast, for parental 786-O cells, serine deprivation alone hindered their invasion capacity, while no significant difference was observed for the combination of sunitinib and NCT-503, irrespective of serine presence in the media (Figure S5B). Overall, our findings suggest that the decrease in cellular serine levels significantly inhibits the motility and invasion of ccRCC cells resistant to sunitinib. However, in sunitinib-sensitive cells, it only inhibits invasion. Unlike the sunitinib-sensitive cells, the presence of serine in the media is crucial for sunitinib-resistant cells, emphasizing the dependence on both intracellular and extracellular sources of serine in modulating invasive and resistance behaviors.

### Sunitinib treatment promotes de novo serine synthesis across diverse cancer models

Given that sunitinib primarily targets Receptor Tyrosine Kinases (RTKs) such as VEGFR2, PDGFRβ, or c-KIT, which are known to be activated in various cancers (^30^), we aimed to broaden our investigation into the effects of sunitinib on serine metabolism across others cancer cell lines. These included BT-549 (triple-negative breast cancer (TNBC)), A549 (lung cancer), DAOY (medulloblastoma), and CAL33 (head and neck cancer). Consistent with our findings in RCC cells, sunitinib treatment led to an increase in protein levels of serine synthesis enzymes (Figure 6A). This indicates that the induction of the serine biosynthetic pathway is not limited to ccRCC but extends to diverse tumor types. Similarly to the strategies employed in ccRCC, we inhibited de novo serine synthesis with the PHGDH inhibitor NCT-503 in these cancer cell lines. Combining sunitinib with NCT-503 resulted in a significant decrease in proliferation after 4 days compared to treatment with sunitinib alone (Figure 6B). These compelling results suggest that the biosynthetic pathway plays a pivotal role in sunitinib resistance, demonstrating its relevance independent of the specific tumor type.

**Figure 6:**
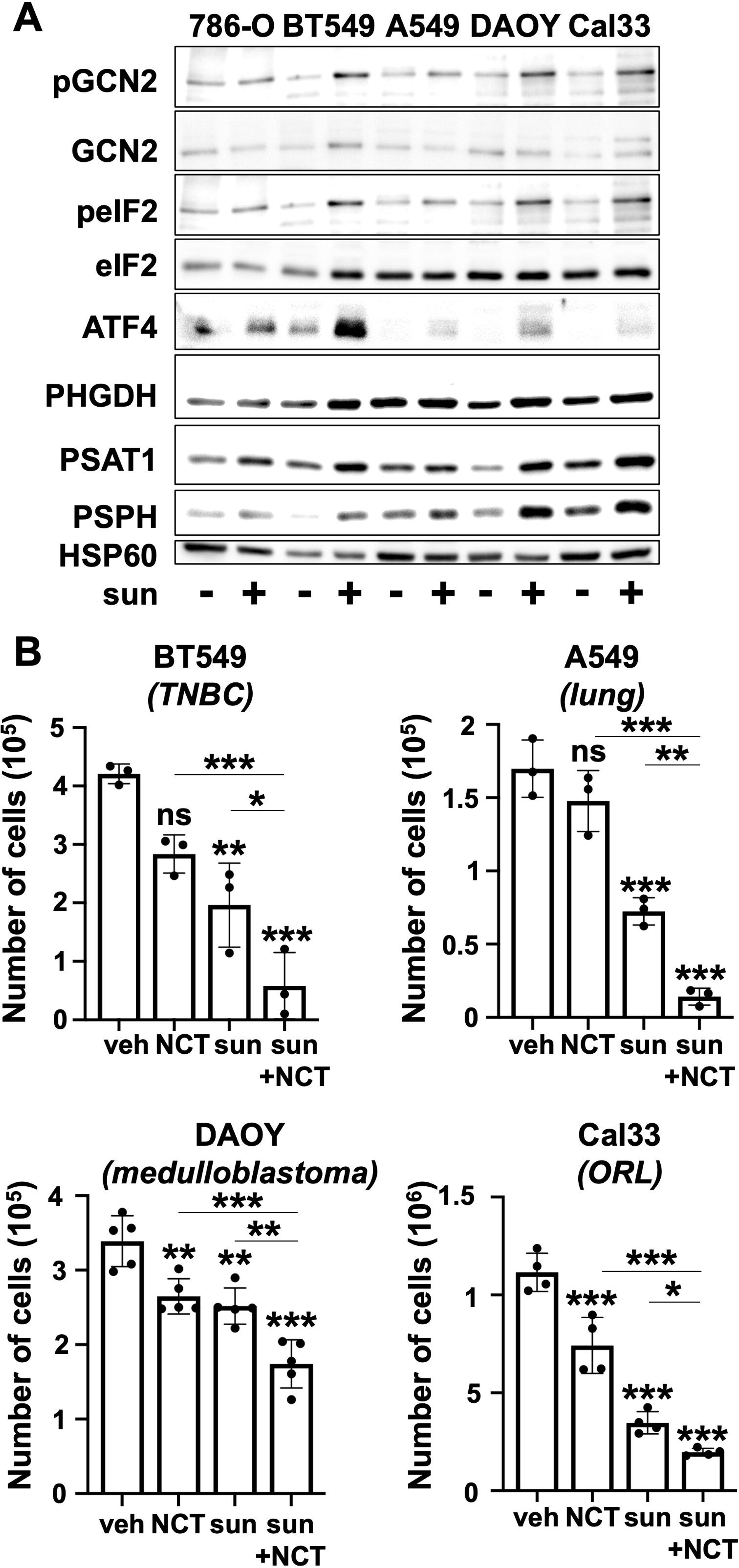
Sunitinib treatment induces an increase in serine de novo metabolism in various models. (A) Protein expression profiles by immunoblots indifferent cancer lines, 786-O (ccRCC), BT549 (TNBC), A549 (lung), DAOY (medulloblastoma), Cal33 (head and neck), treated with DMSO (veh) or sunitinib (sun, 2.5μM) for 48 hours. (B) Cell count measurements conducted 96 hours after treatment of BT549, A549, DAOY, and Cal33 with DMSO (veh), sunitinib (sun), or NCT-503 (NCT), or a combination of both treatments. ns, non-significant; * p < 0.05; ** p < 0.01; *** p < 0.001 (two-way ANOVA).

### Sunitinib treatment promotes serine biosynthesis in RCC tumors and in samples from RCC patients

To investigate whether sunitinib stimulates the de novo serine pathway in tumors, we implanted 786-O cells subcutaneously into nude mice and administered sunitinib (Figure 7A). Metabolite profiling showed distinct metabolic alterations in sunitinib-treated tumors compared to non-treated tumors (Figure S6A-B).

**Figure 7:**
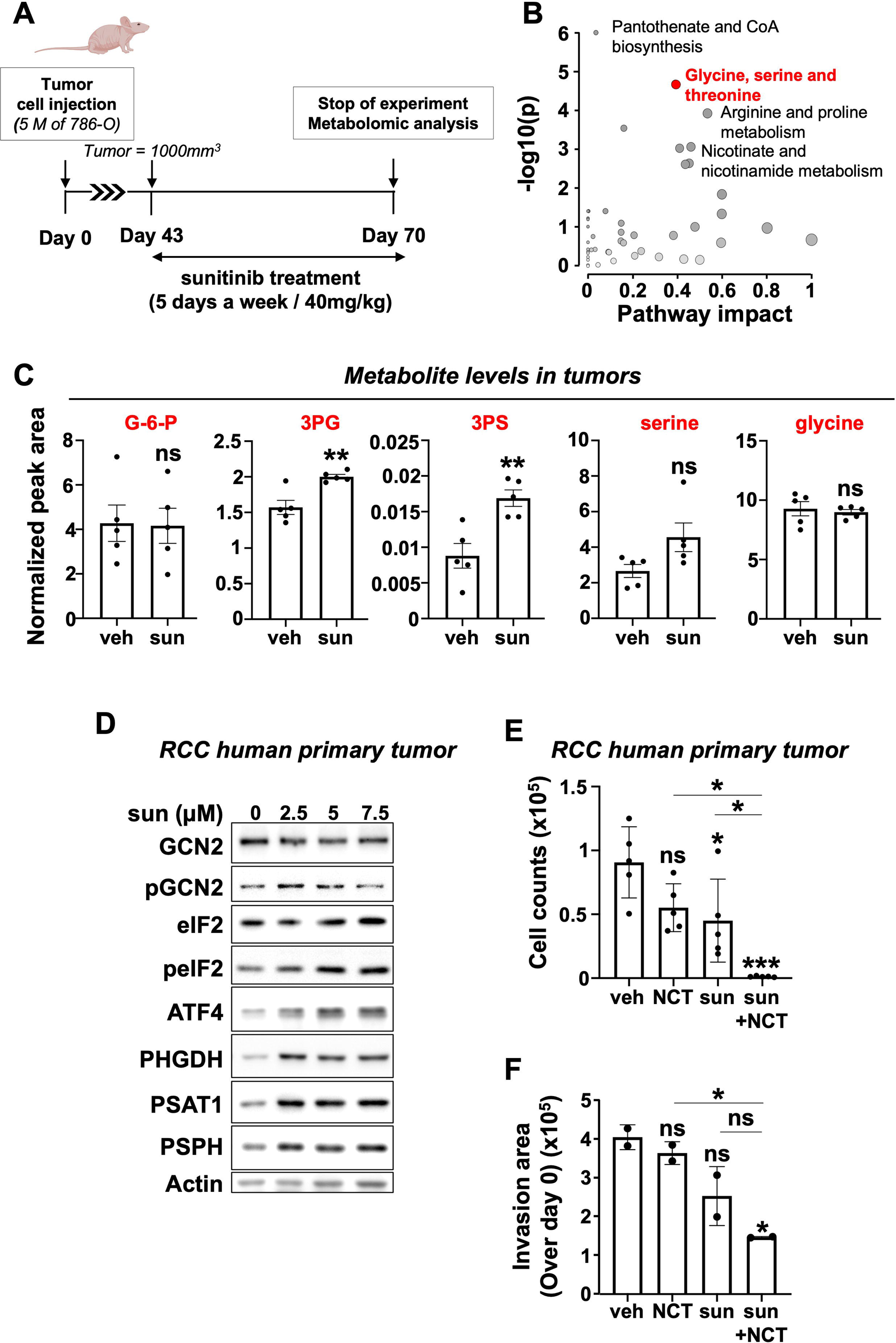
Sunitinib enhances serine biosynthesis pathway in vivo and in ccRCC patient samples. (A) Schematic representation of the in vivo tumor experiment involving subcutaneous implantation of 786-O cells into nude mice (n=5 per group, 2 tumors/mouse). Mice were treated with veh or sunitinib (40 mg/kg/d p.o. d42-d69). In vivo steady-state metabolite profile in RCC xenograft models was shown. (B) Bubble plots of altered metabolic pathways. Bubble plots illustrating altered metabolic pathways in ccRCC xenograft models. Comparison between sunitinib-treated mice and control mice, with bubble area proportional to the effect of each metabolic pathway. The most altered pathways, including glycine, serine, and threonine metabolism are indicated in red (upper right panel). (C) Normalized peak areas of various metabolites intermediates of the serine biosynthesis pathway in ccRCC xenograft models. (D) Immunoblots of fresh human kidney cells obtained by enzymatic dissociation and treated with DMSO (veh) or with sunitinib (sun) at different concentrations for 48 hours. The immunoblot is representative of four different fresh human kidney samples. (E) Cell count measurements performed 96 hours after treatment of fresh human kidney cells with DMSO (veh), sunitinib (sun, 5μM), or NCT-503 (NCT, 50μM), or a combination of both treatments. (F) Quantification of spheroid invasion area after 7 days of invasion. Spheroids performed with cells from cc RCC patient samples were treated with DMSO (veh), sunitinib (sun, 0.5μM), NCT-503 (NCT, 30μM), or a combination of both treatments 24 hours before embedding in Matrigel (1 mg/mL). ns, non-significant; * p < 0.05; *** p < 0.001 (two-way ANOVA).

Significant changes were observed in several major metabolic pathways, including pantothenate and CoA biosynthesis, glycine, serine, and threonine metabolism, and arginine and proline metabolism. Notably, glycine, serine, and threonine metabolism stood out as the second most differentially enriched pathway in RCC tumors from sunitinib-treated mice (Figure 7B, impact factor≥0.1). This was supported by steady-state metabolite profiling, which identified several differentially produced metabolites, including de novo serine synthesis intermediates (Figure 7C).

Consistent with in vitro results, sunitinib-treated tumors exhibited elevated levels of serine and phosphoserine (Figure 7C). Analysis of mRNA expression data from The Cancer Genome Atlas (TCGA) indicated that high expression of *PSAT1* was associated with lower disease-free survival (DFS), progression-free survival (PFS) and overall survival (OS) in renal cancer patients (Figure S7). However, no significant differences were found between renal cancer patients with tumors expressing low or high levels of *PHGDH* and *PSPH* (Figure S7).

Human ccRCC tumor samples obtained from the University Hospital of Nice (CHU Nice) in France, were treated with a sunitinib time course and displayed a rapid increase in PHGDH, PSPH, and PSAT1 protein abundance. Additionally, these RCC tumor cells treated with sunitinib showed an activation of the GCN2-eIF2α axis and an increase in ATF4 protein levels (Figure 7D). These results mirrored findings observed in 786-O cells, supporting the relevance of sunitinib-induced serine synthesis and ISR in ccRCC patients.

Similarly, to observations in 786-O cells, combining NCT-503 with sunitinib remarkably suppressed the proliferation of human RCC tumor cells compared to individual treatments (Figure 7F). Furthermore, PHGDH inhibition notably suppressed the invasiveness of RCC tumor cells (Figure 7G). These findings support the concept that during sunitinib treatment, de novo serine synthesis emerges as a metabolic vulnerability and can be effectively targeted to combat RCC.

### Inactivation of the serine synthesis pathway exhibits anti-tumor and anti-metastasis effects in a zebrafish model

To assess the potential contribution of sunitinib-induced serine synthesis to ccRCC tumor formation and dissemination, we employed a zebrafish model for cancer growth and metastasis studies (PMID: 31615862). Utilizing 786-O cells, known for their efficient metastasis in zebrafish tails (PMID: 29376139), we examined the tumor growth and metastatic potential of ccRCC cells with or without *PSAT1,* treated with either vehicle or sunitinib. 786-O cells, whether depleted of *PSAT1* or not, were labeled with a lipophilic dye (DiI) and injected into the perivitelline space (PVS) of zebrafish embryos at 48 hours post-fertilization (hpf). At 2 days post-injection (dpi), we assessed the extent and growth of the tumor by measuring tumor area and RFP signal intensity (Figure 8A). Quantification analysis revealed a significant reduction in distal metastatic tumor cells when 786-O cells lacked *PSAT1* and zebrafish were treated with sunitinib (Figure 8B-C). Additionally, *PSAT1*-depleted cells exhibited smaller RCC tumors compared to those derived from control ccRCC cells, with further reductions observed during sunitinib treatment (Figure 8D-E). Consistent results were obtained with 786-O cells knockout for *PSAT1* (Figure S8A-B and Figure S9A-D).

**Figure 8:**
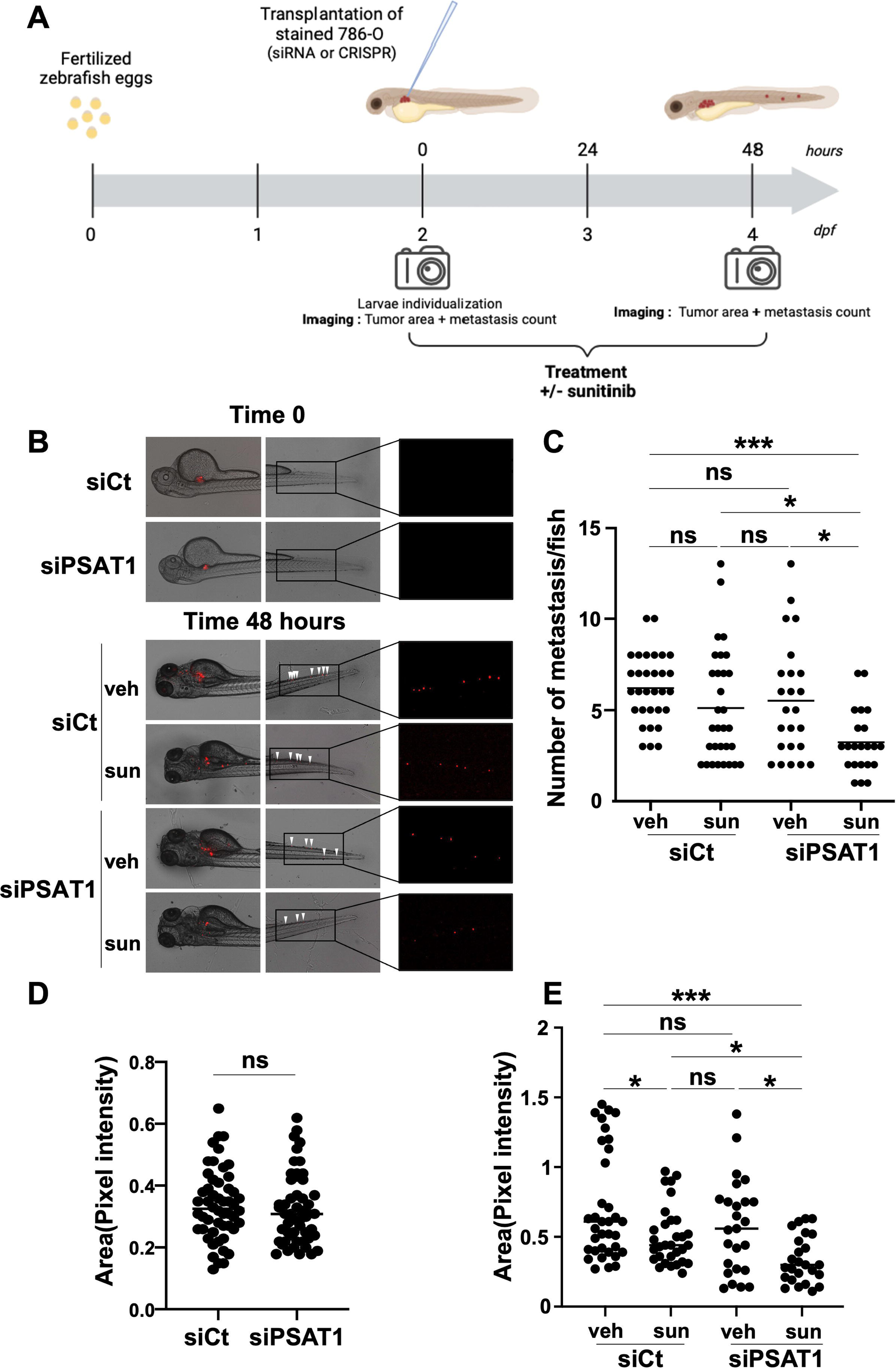
Inhibition of PSAT1 by siRNA suppresses local growth and distant metastases in a zebrafish model. (A) Schematic representation of the in vivo tumor experiment involving the injection of 786-O cells into the perivitelline space of 48 hours post-fertilization zebrafish embryos. (B) Representative images depicting local and distant metastases. Zebrafish embryos (N = 30) were injected with siCt and siPSAT1-treated 786-O cells (labeled with red DiD) into the perivitelline space and treated with sunitinib (2.5 μM). Analysis was conducted at 0 hour and 48 hours later. (C) Zebrafish embryos were monitored for distant tumor metastases using a fluorescent microscope. (D-E) Quantification of tumor growth by measuring tumor area and RFP signal area. ns, non-significant; * p < 0.05; *** p < 0.001 (two-way ANOVA).

In summary, our findings support the role of serine biosynthesis in promoting tumor growth and metastasis. Sunitinib treatment triggers the activation of the GCN2-eIF2α-ATF4 signaling axis, which in turn stimulates the expression of serine synthesis enzymes, subsequently leading to increased de novo serine/glycine synthesis. This metabolic adaptation enables sunitinib-treated cells to withstand stress and persist by promoting nucleotide synthesis, a crucial pathway for sustaining anabolic metabolism and tumor cell survival. Furthermore, inhibition of PHGDH or reduction of PSAT1 levels not only led to decreased cell migration and invasion in metastatic ccRCC cell lines but also demonstrated efficacy in primary RCC cells. These findings reveal a mechanism through which kidney cancer cells persist under sunitinib therapy, involving a metabolic reprogramming that provides essential metabolite building blocks for tumor survival, resistance and progression. Thus, this study could potentially inform future therapeutic strategies targeting serine and nucleotide synthesis pathways to enhance the efficacy of kidney cancer treatments (Graphical abstract).

## DISCUSSION

The intricate landscape of metabolic alterations in response to sunitinib has remained relatively elusive until now. Our study reveals how RCC cells metabolically adapt to sunitinib. Leveraging RNA sequencing and metabolite profiling, we show that RCC cells, sensitive or resistant to sunitinib treatment, exhibit increased serine biosynthesis in sunitinib-treated RCC cells, enabling tumor growth and progression. Indeed, numerous cancer cells, spanning various organs, heavily rely on serine and metabolite production. This unique dependency unveils promising therapeutic prospects, whether through inhibiting de novo serine synthesis or restricting the availability and uptake of exogenous serine.

Crucially, our study extends beyond RCC, revealing that sunitinib treatment enhances serine synthesis in diverse cancer models, including triple-negative breast cancer (TNBC), lung cancer, medulloblastoma, and head and neck cancer. Mechanistically, sunitinib induces the integrated stress response (ISR), activating the GCN2-ATF4 axis, thereby upregulating serine synthesis enzymes and stimulating serine synthesis.

Direct inhibition of serine biosynthesis emerges as a potent strategy, inhibiting tumor growth while curtailing cell migration and invasion. These findings not only deepen our understanding of the metabolic adaptations induced by sunitinib but also offer promising avenues for developing targeted therapies across a spectrum of aggressive cancers. This study was performed on two RCC cell lines, 786-O and RCC10. 786-O which was established as one of the first RCC cell lines, has many characteristics of ccRCC and is most commonly used in RCC research. Like the 786-O RCC line, the RCC10 line is also defective in VHL expression. We also performed an RNAseq analysis in a third RCC line, A498, which also shows a defect in VHL expression. In our model, comprising three distinct cell lines resistant to sunitinib, we observed an elevation in the serine biosynthesis pathway. Interestingly, while PSAT1 and PSPH exhibited an increase, PHGDH showed no significant change.

This led us to hypothesize that PHGDH might be regulated differently compared to PSAT1 and PSPH in our clear cell renal cell carcinoma (ccRCC) model. Specifically, PHGDH regulation may be influenced by HIF2, whereas PSAT1 and PSPH could be modulated by the integrated stress pathway GCN2/eIF2/ATF4 in our ccRCC model (^33;34–36^). Notably, in contrast to our ccRCC model, other cancer cell lines demonstrated an increase in PSAT1, PSPH, and PHGDH with sunitinib treatment, suggesting that the regulation of PHGDH in response to sunitinib may be specific to RCC and unrelated to the treatment itself.

While our findings shed light on these differential expressions, further studies are imperative to enhance our understanding of the intricate pathways governing the upregulation of the serine biosynthesis pathway in ccRCC. ATF4 emerges as a pivotal transcriptional regulator of the serine synthesis pathway in various cancers (^25,26^). In our model, we identified the GCN2 and eIF2 pathway as regulators of ATF4.

Although ATF4 presents a potential target for cancer treatment (^39^), direct ATF4 treatment is hindered by developmental defects observed in Atf4-null mice (^40^). Moreover, siRNA against ATF4 resulted in rapid cell death in our model (data not shown), suggesting that direct targeting of ATF4 may not be feasible.

Recent evidence supports the use of immune checkpoint inhibitors in combination with targeted anti-VEGF agents for treating metastatic renal cell carcinoma. Axitinib, a tyrosine kinase inhibitor similar to sunitinib, did not induce upregulation of the serine biosynthesis pathway in RCC cells, in contrast to sunitinib (data not shown). This discrepancy is attributed to the lysosomotropic properties of sunitinib, which is protonated at acidic pH, preventing its crossing of membranes. Lysosomal sequestration of sunitinib, a process not observed with axitinib, leads to incomplete autophagy, impacting the serine biosynthesis pathway (^8^). Recent studies by Ai-Ling Tian et al. demonstrated that lysosomotropic agents, including azithromycin, chloroquine, and hydroxychloroquine, activate the integrated stress response (^41^). Our initial findings suggest a process specific to lysosomotropic drugs, warranting further investigations to better comprehend the intricate relationship between lysosomotropic drugs and metabolic adaptations. In conclusion, our study unravels the significant role of the serine pathway in inducing resistance across various treatments for RCC and other aggressive tumors. Metabolomic studies offer a novel perspective to identify associated metabolic signatures, revealing that sunitinib treatment or resistance prompts an increase in serine biosynthesis, supporting the growth of treated RCC cells. This phenomenon extends to other cancer models, including TNBC, lung cancer, medulloblastoma, and head and neck cancer.

Mechanistically, sunitinib induces the ISR, activating the GCN2-ATF4 axis, which, in turn, upregulates serine synthesis enzymes and promotes serine synthesis. The direct inhibition of serine biosynthesis not only inhibits tumor growth but also limits cell migration and invasion. Importantly, our findings suggest that targeting the serine biosynthesis pathway could serve as a promising therapeutic strategy to overcome treatment resistance in RCC. Furthermore, our investigation into the impact of lysosomotropic drugs, specifically sunitinib, on the serine biosynthesis pathway reveals a unique relationship. Unlike other tyrosine kinase inhibitors, sunitinib’s lysosomal sequestration induces incomplete autophagy, affecting ATF4 expression and subsequently influencing serine biosynthesis. This intriguing link between lysosomotropic drugs and metabolic adaptations merits further exploration to better comprehend its implications in cancer treatment.

In essence, our study sheds light on the dynamic interplay between sunitinib treatment, the serine biosynthesis pathway, and the integrated stress response. These insights not only enhance our understanding of the metabolic mechanisms facilitating resistance to current therapies used against ccRCC, but also unveil promising therapeutic strategies to counter resistance and tackle aggressive RCC by targeting the serine biosynthesis pathway.

## SUPPLEMENTAL INFORMATION

Supplemental information can be found online at https://doi.org/10.1016/j.cmet.2023.12.014.

## ACKNOWLEDGMENTS

This study was supported by the French Ministry of research (PhD grant MT), This work was supported by funding from the Fondation de France GP, the Fondation ARC “Equipe Labellisée” and the contract ARCAGEING2023020006332 GP and PJA20191209349 SG, the national agency of research France (ANR, JCJC contract, ANR-21-CE14-0008-01, SG), the Ligue Nationale contre le Cancer Equipe labellisée 2019 GP, Canceropole PACA (Emergence, 2019, SG), UCA credits incitatifs SG, and the National Institutes of Health (NIH), R01GM135587, R01GM143334-04 (IB-S).

## AUTHOR CONTRIBUTIONS

Conceptualization, IB-S, SG; methodology, MT, US, YC; investigation, MT, US, MC-T, JP, VV, JD, RG, YC; writing– original draft GP, IB-S, SG; writing – review and editing, all authors; funding acquisition, GP, IB-S, SG; resources, GP, IB-S, SG; project administration, GP, IB-S, SG; and supervision GP, IB-S, SG.

## DECLARATION OF INTERESTS

The authors declare no competing interests.

## INCLUSION AND DIVERSITY

We support inclusive, diverse, and equitable conduct of research.

## Supplemental information

**Supplemental Figure 1:**
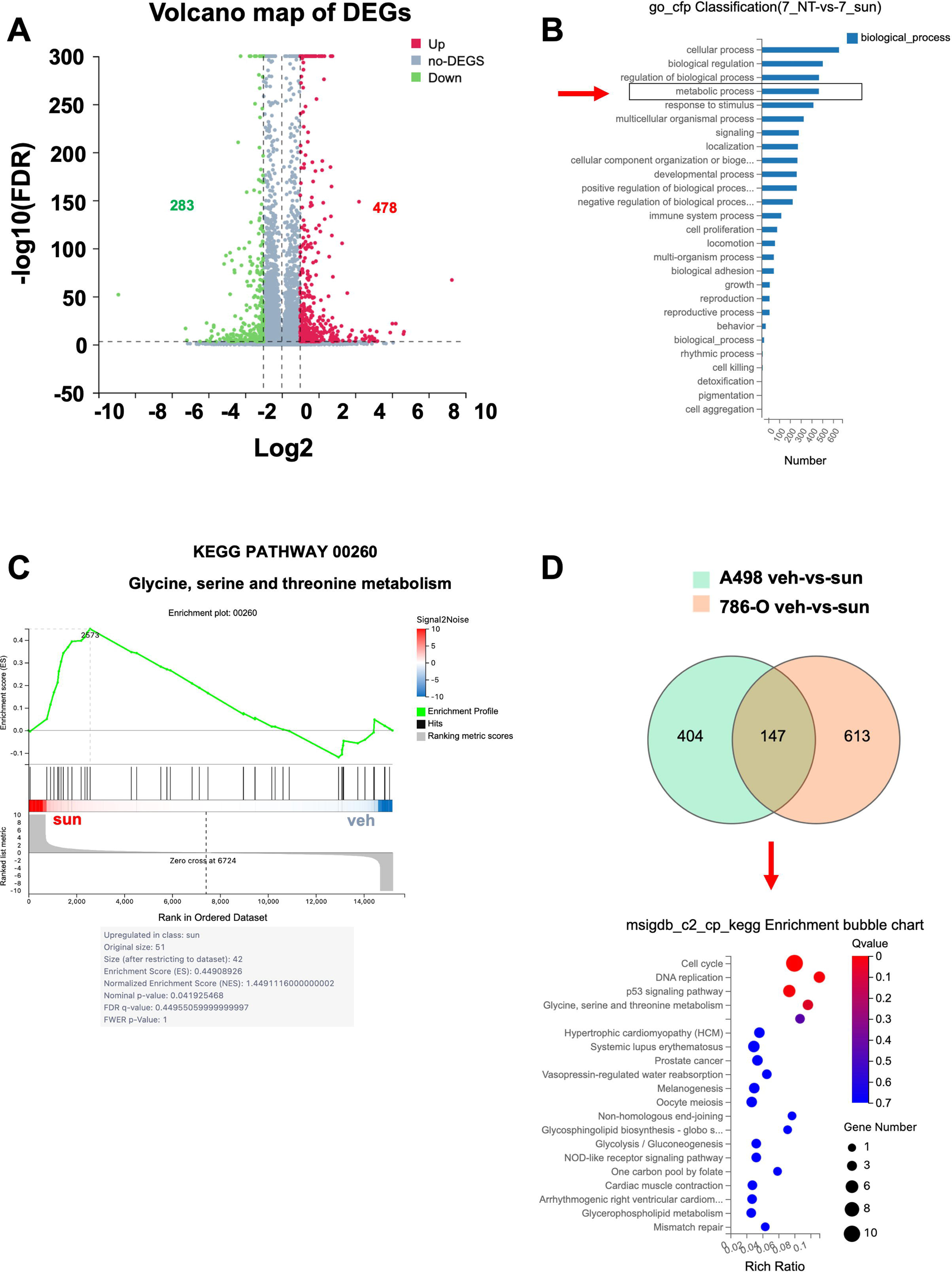
Modulated Signaling Pathways in Sunitinib-Treated and Resistant Cells. (A) RNA-seq analysis revealed 761 genes with abnormal regulation with p < 0.05 and fold change (sun/C) > 2 or the reciprocal value of fold change > 2. C, control; sun, sunitinib. (B) Functional analyses of RNA-seq results were conducted using Gene Ontology (GO) terms. Within the Biological Process category, genes clustered into 27 GO terms, with “metabolic process” ranking among the top five terms. (C) Using the KEGG pathway computational tool, functional analyses of RNA-seq results revealed significant enrichment in the “glycine, serine and threonine metabolism” pathway in 786-O parental cells treated with sunitinib (sun) compared to cells treated with DMSO (veh). (D) Venn diagram illustrating the overlap of significant transcripts between 786-O and A498 cells treated with suninitib based on RNA-seq data analysis. Functional enrichment results based on KEGG, are presented as dot-bubble diagrams, with pathways arranged in ascending order of significance from top to bottom.

**Supplemental Figure 2:**
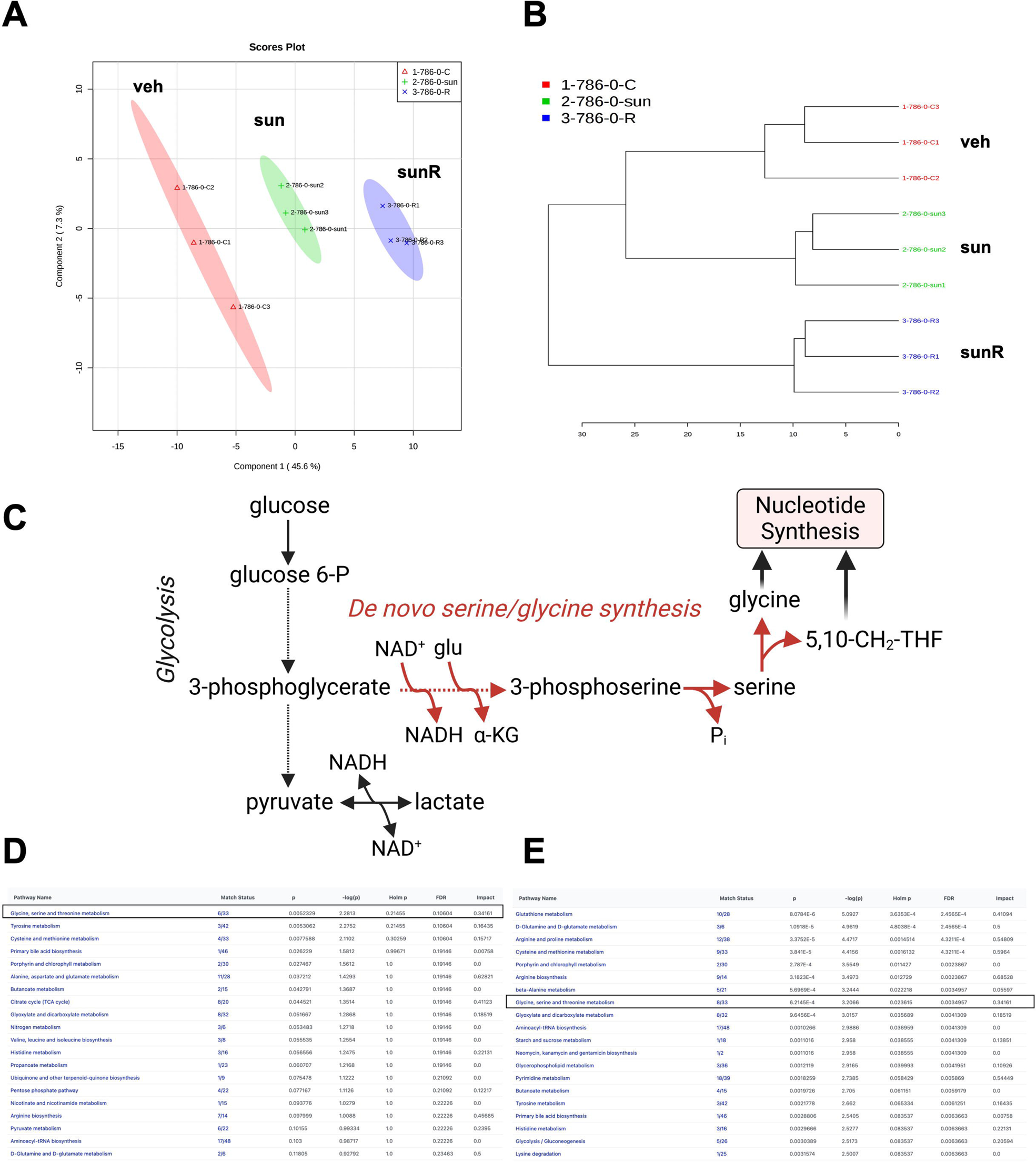
Molecular Responses to Sunitinib Treatment and Resistance. (A-B) Dendrogram illustrating the steady-state metabolomic analysis conducted in triplicate for of three conditions: control (veh), sunitinib-treated cells (sun), and cells resistant to sunitinib (sunR). The vertical axis represents the complexes analyzed as samples, while the horizontal axis indicates the average distance between different clusters. (C) Schematic representation of the de novo serine synthesis pathway of serine. **(D-E)** Table listing the top 20 major signaling pathways identified as modulated in sunitinib-treated cells (left) or in sunitinib-resistant cells (right). The pathways are ordered from the highest to lowest modulation determined using the Pathway Analysis module of Metaboanalyst 5.0.

**Supplemental Figure 3:**
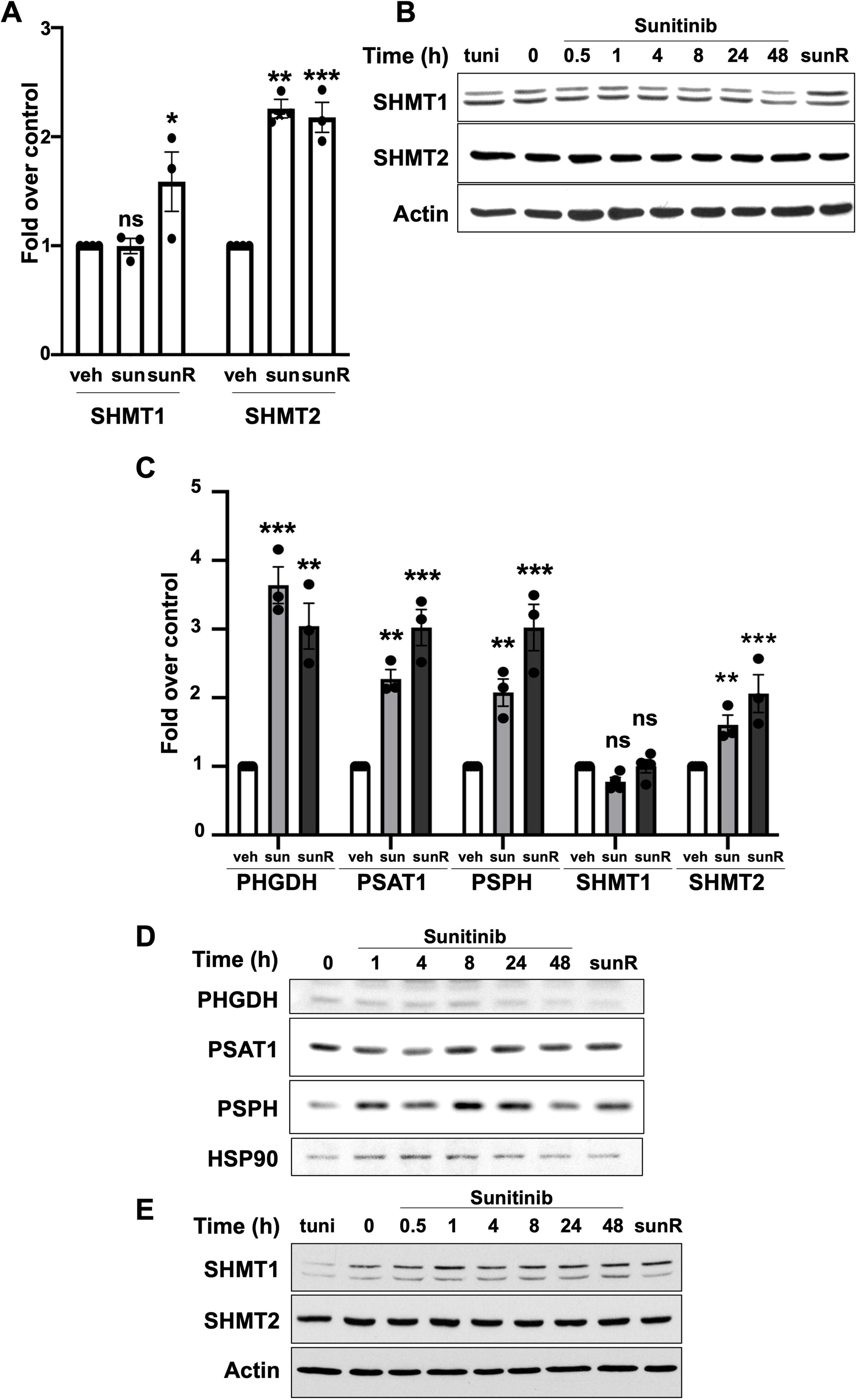
Molecular and functional responses to sunitinib treatment and resistance. (A) mRNA levels of SHMT1 and SHMT2 in 786-O cells treated with sunitinib (sun, 2.5μM, 48h) or in cells resistant to this treatment (sunR). (B) Immunoblots of 786-O Parental cells treated with sunitinib (sun, 2.5μM) at different time points or in cells resistant to this treatment (sunR). Tunicamycin (tuni, 48h, 1μg/ml) serves as a control. (C) mRNA levels of PHGDH, PSAT1, and PSPH in RCC10 cells treated with sunitinib (sun, 2.5μM, 48h) or from cells resistant to this treatment (sunR). (D) Immunoblots of RCC10 Parental cells treated with sunitinib at different time points or from cells resistant to this treatment (sunR). (E) Immunoblots of RCC10 Parental cells treated with sunitinib (sun, 2.5μM) at different time points or of cells resistant to this treatment (sunR). Tunicamycin (tuni, 48h, 1μg/ml) is used as a control. ns, non-significant; * p < 0.05; *** p < 0.001 (two-way ANOVA).

**Supplemental Figure 4:**
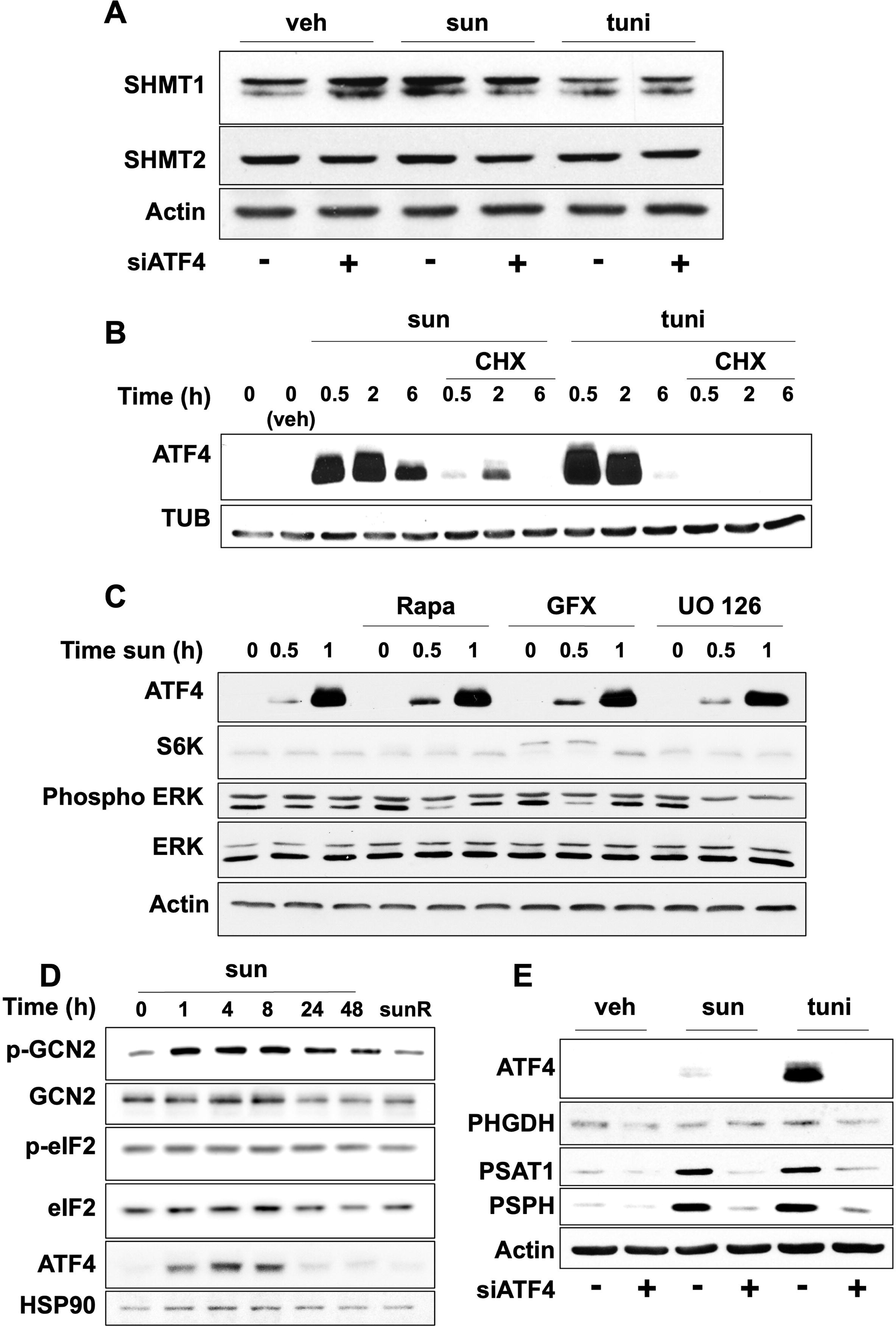
Protein stability and inhibitor effects. (A) Immunoblots of 786-O cells treated with DMSO (veh), sunitinib (sun, 2.5μM) or tunicamycin (tuni, 1μg/ml) and transfected with siRNAs targeting ATF4 (siATF4) or non-targeting controls for 48 hours before protein extraction. (B) Immunoblots of 786-O cells treated with DMSO (veh) or with a time course of sunitinib (sun, 2.5μM) or tunicamycin (tuni, 1μg/ml) in the presence or absence of the translational inhibitor cycloheximide (CHX, 10μM). (C) Immunoblots of 786-O cells treated with DMSO (veh) or with a time course of sunitinib (sun, 2.5μM) in the presence or absence of various inhibitors added 1 hour before sunitinib treatment. Inhibitors include rapamycin (rapa, 20nM), GF109203X (GFX, 5μM), or UO126 (UO, 10μM). (D) Immunoblots of RCC10 Parental cells treated or not with sunitinib at different time points or from cells resistant to this treatment (sunR). (E) Immunoblots of 786-O cells treated with DMSO (veh), sunitinib (sun, 2.5μM) or tunicamycin (tuni, 1μg/ml) and transfected with siRNAs targeting ATF4 (siATF4) or non-targeting controls for 48 hours before protein extraction.

**Supplemental Figure 5:**
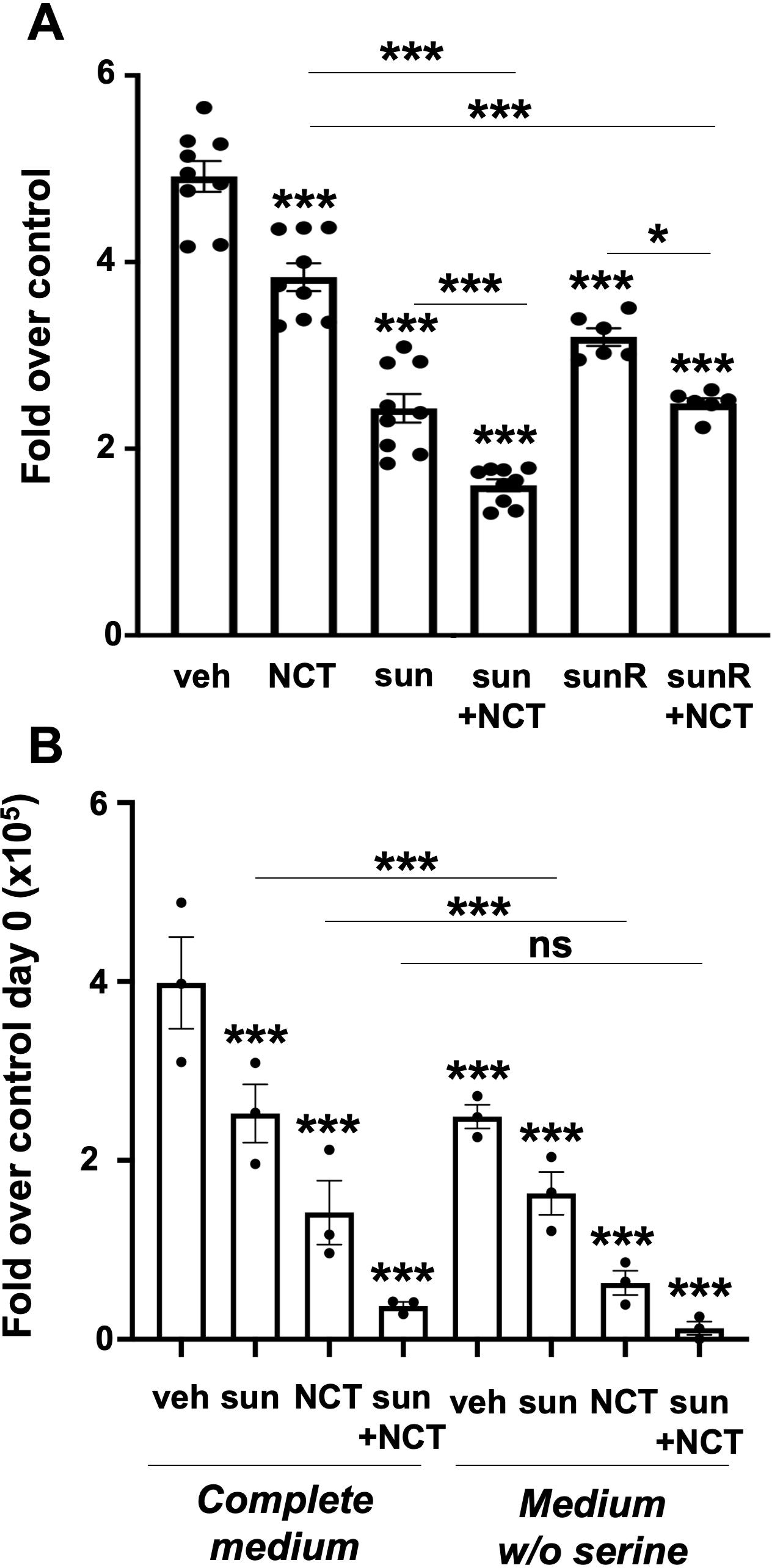
Impact of combined treatments targeting the serine pathway with sunitinib, as well as the effects of culturing ccRCC cells in serine-containing and serine-free media, in both 2D and 3D (spheroid) culture models. (A) Cell Titer-Glo measurements performed 96 hours after treatment of RCC10 cells with DMSO (veh), sunitinib (sun, 2.5μM), NCT-503 (NCT, 30μM), or a combination of both treatments. (B) Spheroids of 786-O parental cells were treated with DMSO (veh), sunitinib (sun, 2.5μM), NCT-503 (NCT, 30μM), or a combination of both treatments 24 hours before embedding in Matrigel (1 mg/mL) and cultured in serine-containing (complete media) or serine-free media (media w/o serine). ns, non-significant; ** p < 0.01; *** p < 0.001 (two-way ANOVA).

**Supplemental Figure 6:**
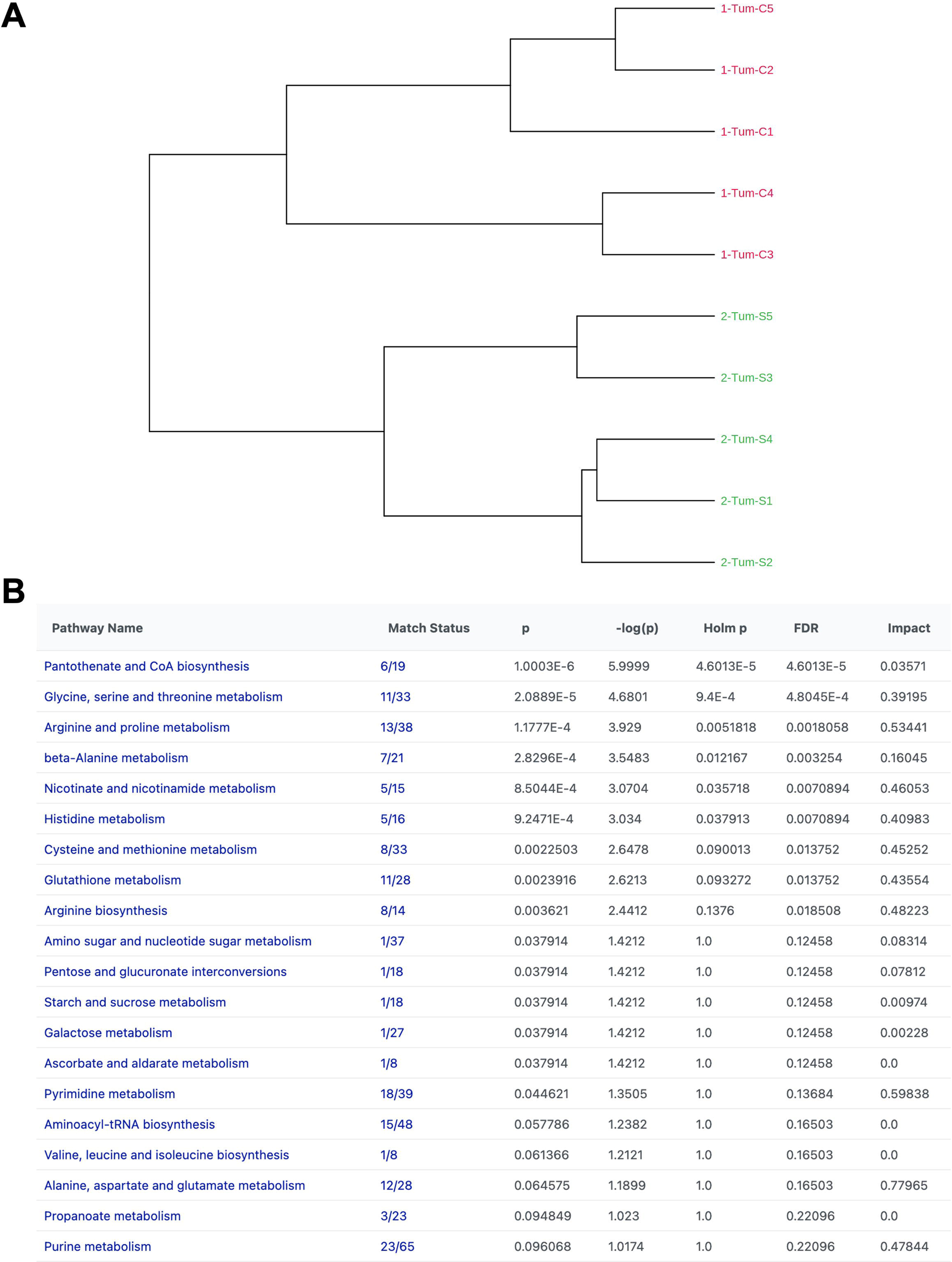
Xenograft Model and Metabolomic Analysis. Xenograft model after subcutaneous injection of 5 × 10^6^ 786-O in nude mice. (A) Dendrogram illustrating the steady-state metabolomic analysis of two control conditions (C) and sunitinib-treated cells (sun) in five mice per group. The vertical axis represents the complexes analyzed as samples and the horizontal axis represents the average distance between different clusters. (B) List of metabolic pathways generated by MetaboAnalyst 5.0. ns, non-significant; * p < 0.05 (two-way ANOVA).

**Supplemental Figure 7:**
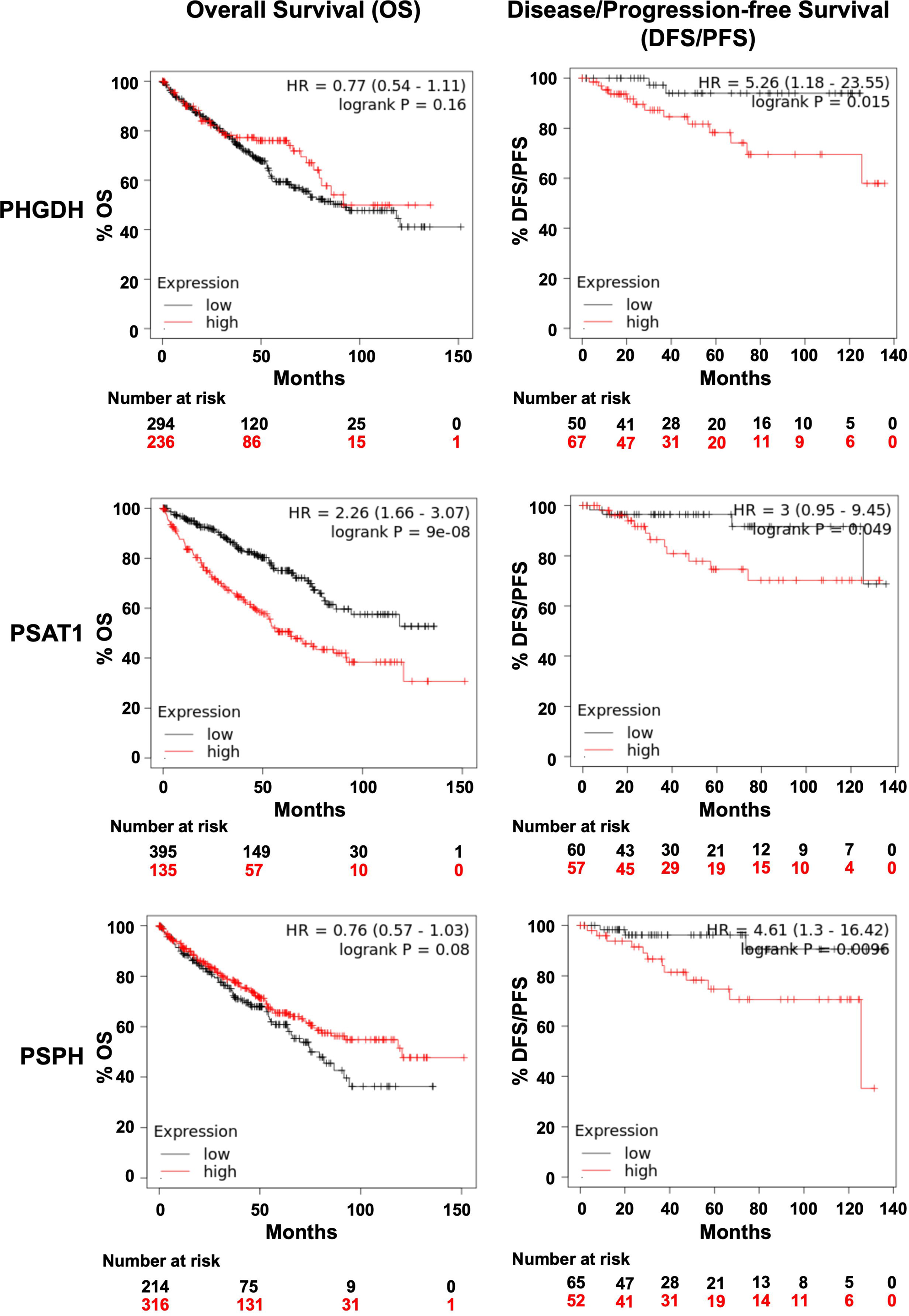
RCC patient outcome depended on the expression of genes of the de novo serine synthesis. 535 patients from The Cancer Genome Atlas (TCGA) were analyzed for mRNA levels of the PHGDH, PSAT1 and PSPH genes and subsequent follow-up in terms of disease-free survival (DFS), progression-free survival (PFS) and overall survival (OS).

**Supplemental Figure 8:**
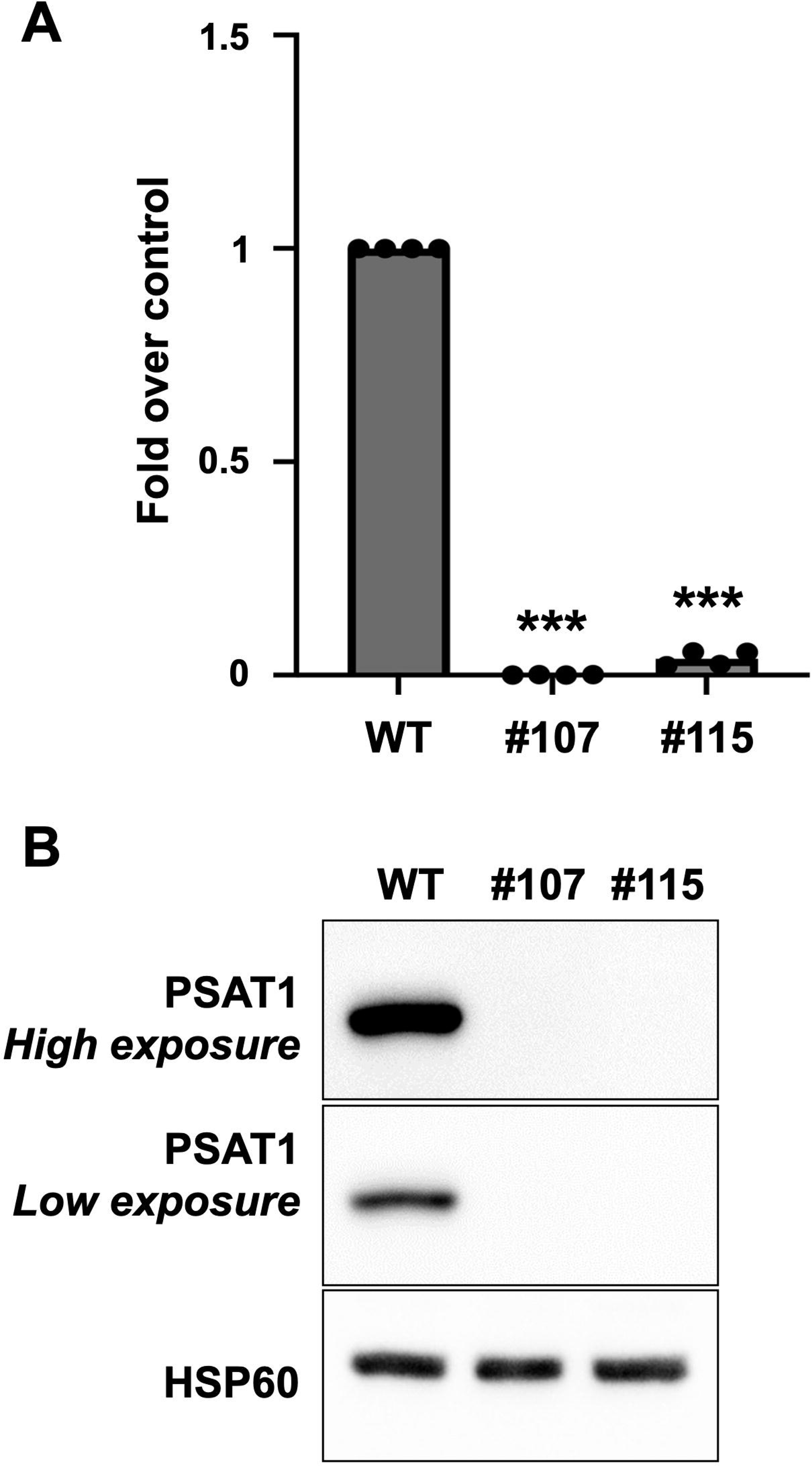
PSAT1 invalidation by CRISPR-Cas9 technology. Invalidation of PSAT1 by CRISPR-Cas9 technology. Clones #107 and #115 were selected for subsequent experiments. (A) The mRNA level of PSAT1 was monitored by RT-QPCR (B) The expression of PSAT1 was also monitored by immunoblot in the selected clones. *** p < 0.001 (two-way ANOVA).

**Supplemental Figure 9:**
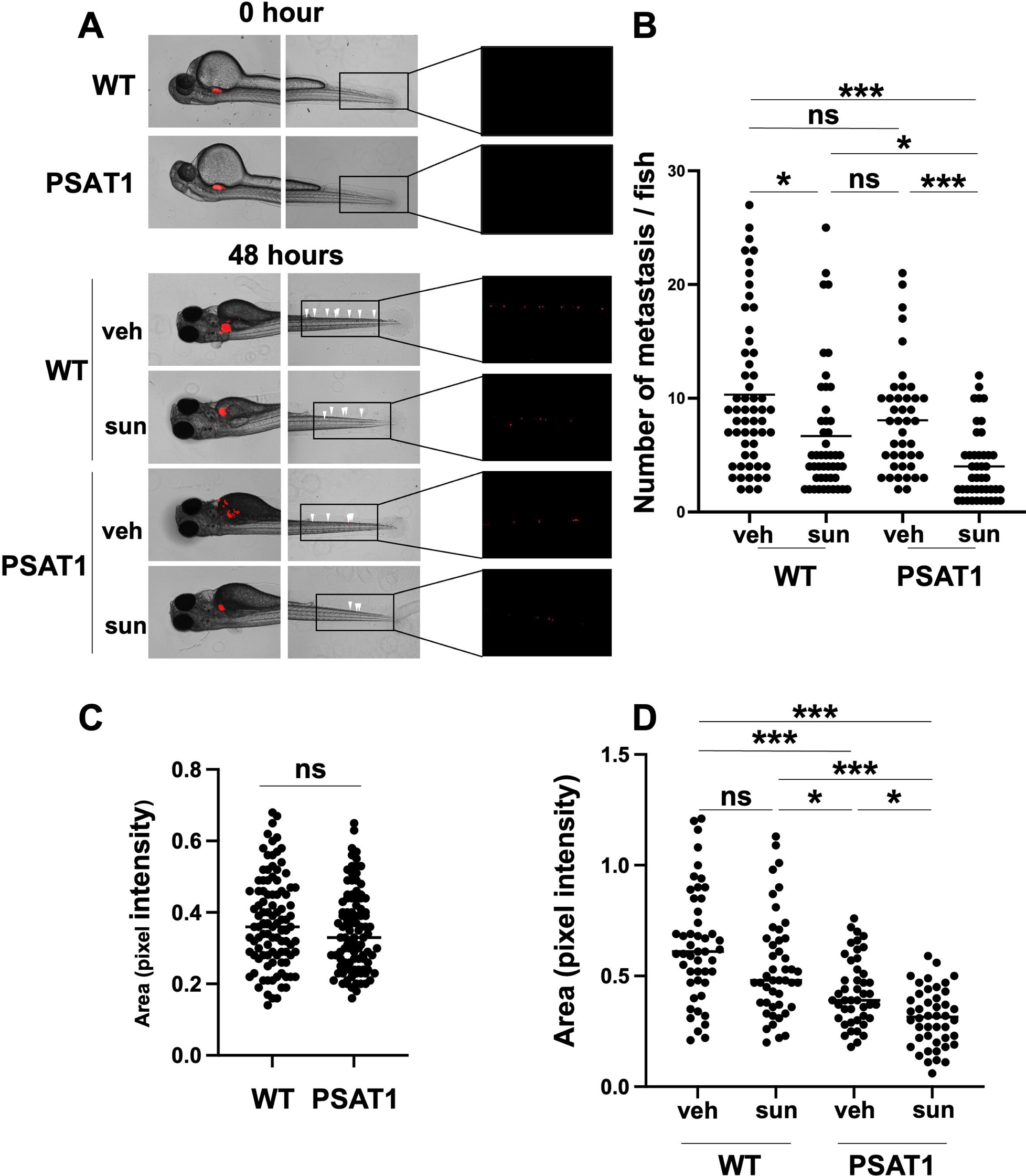
In vivo zebrafish tumor experiment with CRISPR PSAT1 cells. (A) Representative images depicting local and distant metastases. Zebrafish embryos (N = 30) were injected with CRISPR WT and CRISPR PSAT1 786-O cells (labeled with red DiD) into the perivitelline space and treated with sunitinib (2.5 μM) and analyzed at 0 hour and 48 hours later. (B) Zebrafish embryos were monitored for distant tumor metastases using a fluorescent microscope. (C-D) Quantification of tumor growth by measuring tumor area and RFP signal area. ns, non-significant; * p < 0.05; *** p < 0.001 (two-way ANOVA).

## Notes

### Competing Interest Statement

The authors have declared no competing interest.

## References

1. Maxwell, P. H. et al. The tumour suppressor protein VHL targets hypoxia-inducible factors for oxygen-dependent proteolysis. Nature 399, 271–275 (1999).

2. Kaelin, W. G. Molecular basis of the VHL hereditary cancer syndrome. Nat. Rev. Cancer 2, 673–682 (2002).

3. Brugarolas, J. PBRM1 and BAP1 as Novel Targets for Renal Cell Carcinoma. Cancer J. Sudbury Mass 19, 324–332 (2013).

4. Escudier, B. et al. Renal cell carcinoma: ESMO Clinical Practice Guidelines for diagnosis, treatment and follow-up†. Ann. Oncol. Off. J. Eur. Soc. Med. Oncol. 30, 706–720 (2019).

5. Kollmannsberger, C. et al. Sunitinib therapy for metastatic renal cell carcinoma: recommendations for management of side effects. Can. Urol. Assoc. J. 1, S41–S54 (2007).

6. Sehdev, S. Sunitinib toxicity management – a practical approach. Can. Urol. Assoc. J. 10, S248–S251 (2016).

7. Motzer, R. J. et al. Overall Survival and Updated Results for Sunitinib Compared With Interferon Alfa in Patients With Metastatic Renal Cell Carcinoma. J. Clin. Oncol. 27, 3584–3590 (2009).

8. Giuliano, S. et al. Resistance to sunitinib in renal clear cell carcinoma results from sequestration in lysosomes and inhibition of the autophagic flux. Autophagy 11, 1891–1904 (2015).

9. Reid, M. A. et al. Serine synthesis through PHGDH coordinates nucleotide levels by maintaining central carbon metabolism. Nat. Commun. 9, 5442 (2018).

10. Fan, T. W. M. et al. De novo synthesis of serine and glycine fuels purine nucleotide biosynthesis in human lung cancer tissues. J. Biol. Chem. 294, 13464–13477 (2019).

11. DeBerardinis, R. J. Serine metabolism: some tumors take the road less traveled. Cell Metab. 14, 285–286 (2011).

12. Pan, S., Fan, M., Liu, Z., Li, X. & Wang, H. Serine, glycine and one-carbon metabolism in cancer (Review). Int. J. Oncol. 58, 158–170 (2020).

13. Pacold, M. E. et al. A PHGDH inhibitor reveals coordination of serine synthesis and 1-carbon unit fate. Nat. Chem. Biol. 12, 452–458 (2016).

14. Ou, Y., Wang, S.-J., Jiang, L., Zheng, B. & Gu, W. p53 Protein-mediated Regulation of Phosphoglycerate Dehydrogenase (PHGDH) Is Crucial for the Apoptotic Response upon Serine Starvation. J. Biol. Chem. 290, 457–466 (2015).

15. Frezza, C. Cancer metabolism: Addicted to serine. Nat. Chem. Biol. 12, 389–390 (2016).

16. Labuschagne, C. F., van den Broek, N. J. F., Mackay, G. M., Vousden, K. H. & Maddocks, O. D. K. Serine, but not glycine, supports one-carbon metabolism and proliferation of cancer cells. Cell Rep. 7, 1248–1258 (2014).

17. Locasale, J. W. et al. Phosphoglycerate dehydrogenase diverts glycolytic flux and contributes to oncogenesis. Nat. Genet. 43, 869–874 (2011).

18. Possemato, R. et al. Functional genomics reveals serine synthesis is essential in PHGDH-amplified breast cancer. Nature 476, 346–350 (2011).

19. Mullarky, E., Mattaini, K. R., Vander Heiden, M. G., Cantley, L. C. & Locasale, J. W. PHGDH amplification and altered glucose metabolism in human melanoma. Pigment Cell Melanoma Res. 24, 1112–1115 (2011).

20. Rathore, R., Schutt, C. R. & Van Tine, B. A. PHGDH as a mechanism for resistance in metabolically-driven cancers. Cancer Drug Resist. 3, 762–774 (2020).

21. Masson, G. R. Towards a model of GCN2 activation. Biochem. Soc. Trans. 47, 1481–1488 (2019).

22. Melatonin protects against environmental stress-induced fetal growth restriction via suppressing ROS-mediated GCN2/ATF4/BNIP3-dependent mitophagy in placental trophoblasts - ScienceDirect. https://www.sciencedirect.com/science/article/pii/S2213231721000021.

23. Gao, L. et al. Microcystin-LR inhibits testosterone synthesis via reactive oxygen species-mediated GCN2/eIF2α pathway in mouse testes. Sci. Total Environ. 781, 146730 (2021).

24. Harding, H. P. et al. Regulated translation initiation controls stress-induced gene expression in mammalian cells. Mol. Cell 6, 1099–1108 (2000).

25. Harding, H. P. et al. An Integrated Stress Response Regulates Amino Acid Metabolism and Resistance to Oxidative Stress. Mol. Cell 11, 619–633 (2003).

26. DeNicola, G. M. et al. NRF2 regulates serine biosynthesis in non-small cell lung cancer. Nat. Genet. 47, 1475–1481 (2015).

27. Ye, J. et al. Pyruvate kinase M2 promotes de novo serine synthesis to sustain mTORC1 activity and cell proliferation. Proc. Natl. Acad. Sci. U. S. A. 109, 6904–6909 (2012).

28. Scheuner, D., et al. Translational Control Is Required for the Unfolded Protein Response and In Vivo Glucose Homeostasis. Mol. Cell 7, 1165–1176 (2001).

29. Lu, P. D., Harding, H. P. & Ron, D. Translation reinitiation at alternative open reading frames regulates gene expression in an integrated stress response. J. Cell Biol. 167, 27–33 (2004).

30. Park, Y., Reyna-Neyra, A., Philippe, L. & Thoreen, C. C. mTORC1 Balances Cellular Amino Acid Supply with Demand for Protein Synthesis through Post-transcriptional Control of ATF4. Cell Rep. 19, 1083–1090 (2017).

31. Metcalf, S. et al. Selective loss of phosphoserine aminotransferase 1 (PSAT1) suppresses migration, invasion, and experimental metastasis in triple negative breast cancer. Clin. Exp. Metastasis 37, 187–197 (2020).

32. Zhu, S., Wang, X., Liu, L. & Ren, G. Stabilization of Notch1 and β-catenin in response to ER-breast cancer-specific up-regulation of PSAT1 mediates distant metastasis. Transl. Oncol. 20, 101399 (2022).

33. Huang, D. et al. Sunitinib Acts Primarily on Tumor Endothelium rather than Tumor Cells to Inhibit the Growth of Renal Cell Carcinoma. Cancer Res. 70, 1053–1062 (2010).

34. Yoshino, H. et al. PHGDH as a key enzyme for serine biosynthesis in HIF2α-targeting therapy for renal cell carcinoma. Cancer Res. 77, 6321–6329 (2017).

35. Samanta, D. et al. PHGDH Expression Is Required for Mitochondrial Redox Homeostasis, Breast Cancer Stem Cell Maintenance, and Lung Metastasis. Cancer Res. 76, 4430–4442 (2016).

36. Samanta, D., Prabhakar, N. R. & Semenza, G. L. Systems Biology of Oxygen Homeostasis. Wiley Interdiscip. Rev. Syst. Biol. Med. 9, 10.1002/wsbm.1382 (2017).

37. Semenza, G. L. Hypoxia-inducible factors: coupling glucose metabolism and redox regulation with induction of the breast cancer stem cell phenotype. EMBO J. 36, 252–259 (2017).

38. Yang, M. & Vousden, K. H. Serine and one-carbon metabolism in cancer. Nat. Rev. Cancer 16, 650–662 (2016).

39. Masuoka, H. C. & Townes, T. M. Targeted disruption of the activating transcription factor 4 gene results in severe fetal anemia in mice. Blood 99, 736–745 (2002).

40. Tian, A.-L. et al. Lysosomotropic agents including azithromycin, chloroquine and hydroxychloroquine activate the integrated stress response. Cell Death Dis. 12, 1–13 (2021).

