## Supplementary material for "Exploiting De Novo Serine Synthesis as a Metabolic Vulnerability to Overcome Sunitinib Resistance in Advanced Renal Cell Carcinoma": Material & Methods

### Lead contact

### Materials availability

This study did not generate new unique reagents.

### Data and code availability

This paper does not report original code. Any additional information required to reanalyze the data reported in this paper is available from the lead contact upon request.

**EXPERIMENTAL MODEL DETAILS**

Animals

**METHOD DETAILS**

Cell culture

Cell count and viability

Cell Titer-Glo

Immunoblotting

Quantitative Real-Time PCR (qPCR) experiments

Colony Formation Assay

Migration assay/ Scratch assay

Cell migration/Boyden chambers

Spheroid formation and invasion assay

Metabolite profiling for targeted steady state and flux analyses

Ectopic model of RCC

siRNA assay

Enzymatic dissociation of tumor tissues

Gene expression microarray analysis

Measurements of glucose, glycine, hypoxanthine, serine, and formate uptakes

CRISPRCas9 generation

QUANTIFICATION AND STATISTICAL ANALYSIS

**Materials**

Sunitinib came from residual materials given to patients (Centre Antoine Lacassagne Nice France) and prepared as a 2.5 mmol/L stock solution in dimethyl sulfoxide (DMSO; Sigma-Aldrich) and stored at -20°C. Chloroquine (C6628) was purchased from Sigma Aldrich and NCT-503 from MedChemTronica (HY-101966).

Anti-PHGDH (14719-1-AP), Anti-PSPH (14513-1-AP), Anti-SHMT1 (14149-1-AP), Anti-SHMT2 (11099-1-AP), Anti-Actin (66009-1-Ig), Anti-Beta-Tubulin (66031-1-Ig) were obtained from Proteintech. Anti-GCN2 (#3302), anti-peIF2 (#3398), anti-ATF4 (#11815), anti-S6K (#9202), anti-HSP60 (#12165), and anti-HSP90 (#4877) were purchased from Cell Signaling Technology. Anti-pGCN2 (#75836) and anti-eIF2 (#5369) were obtained from Abcam. Anti-PSAT1 was obtained from Invitrogen (PA5-22124).

**Cell culture**

Human RCC 786-O cell line (CRL-1932) and A498 cell line (HTB-44) were purchased from the American Tissue Culture Collection (ATCC) and RCC10 were a kind gift from W.H. Kaelin (Dana-Farber Cancer Institute, Boston, MA). RCC cells were grown in DMEM high glucose, GlutaMAXTM Supplement, pyruvate and supplemented with 7% FBS. Resistant cells were obtained by a chronic exposition to increasing concentrations of sunitinib up to 8 μmoL/L. Primary human RCC cells (isolated by enzymatic dissociation from the exeresis of the operative specimen obtained thanks to Dr D. Ambrosetti, CHU Nice, department of pathology) were grown in PromoCell Renal Epithelial Cell Growth Medium 2 (PromoCell).

CAL33 HNSCC cells, a human head and neck cancer cell line, was purchased from DSMZ (HTB-186). Cells were cultured in DMEM high glucose, GlutaMAXTM Supplement, pyruvate and supplemented with 7% FBS.

Human medulloblastoma cell line Daoy (HTB-186) was purchased from ATCC and maintained in MEMα with Glutamax (Thermo Fisher, Montigny-le-Bretonneux, France) supplemented with 10% FBS and 1 mM sodium pyruvate.

The TNBC cell line, BT-549 (HTB-122) was obtained from the ATCC and was cultured with DMEM high glucose, GlutaMAXTM Supplement, pyruvate supplemented with 10% of FBS and 1% of non-essential Amino acid.

A549 cells are lung carcinoma epithelial cells obtained from ATCC (CRM-CCL-185) and grown in DMEM high glucose, GlutaMAXTM Supplement, pyruvate and supplemented with 7% FBS.

All cells were grown at 37°C in a humidified atmosphere containing 5% CO2.

**Cell count and viability**

Cells were seeded in 12-well plates at density of 30 000 cells (48h) or 15 000 cells (96h) per well in media containing 7% FBS (fetal bovine serum) and treated with vehicule (DMSO) or sunitinib and/or NCT-503. Cell count and cell viability were assessed using the ADAM-MC apparatus (NanoEnTek) based on fluorescent PropidiumIodide (PI) staining according to the manufacturer’s instructions. All cell count and viability assays were performed in biological triplicates seeded in technical triplicates for each condition.

**Cell Titer-Glo**

Cells were seeded in 96-well plates at density of 1000 cells per well in media containing 7% FBS treated with vehicule (DMSO) or sunitinib (sun, 2.5 µM) and/or NCT-503 (NCT, 30 µM). Cell proliferation was measured using CellTiter-Glo® at 96 hours. The data were normalized to day 0. All cell proliferation assays were performed in biological triplicates seeded in technical triplicates for each condition.

**Immunoblotting**

To extract proteins, cells were lysed in ice-cold Triton lysis buffer (40 mM HEPES, pH 7.4, 120 mM NaCl, 1 mM EDTA, 1% Triton X-100, 10 mM glycerol 2-phosphate, 10 mM sodium pyrophosphate, 0.5 mM sodium orthovanadate and 50 mM NaF; 1 µM Microcystin-LR and protease inhibitor cocktail were added prior to the lysis of cells) and incubated on ice for 30 min. Lysates were centrifuged at 20,000 × g for 15 min at 4°C. Protein concentrations were quantified by BCA assay and protein lysates were normalized for each experiment. Identical amounts of protein lysates (~15 to 20 µg) were prepared with the loading buffer (Laemmli buffer), incubated at 95°C on heat block for 5 min and separated on SDS-PAGE and transferred onto nitrocellulose blotting membranes. Membranes were blocked in 5% milk in TBST (50 mM Tris pH 7.4, 150 mM NaCl, 0.1% Tween 20) and incubated overnight at 4°C with the indicated primary antibodies followed by incubation with horseradish peroxidase (HRP)-tagged anti-rabbit or anti-mouse secondary antibodies for one hour. Proteins were then visualized with the ECL system.

**Quantitative Real-Time PCR (qPCR) experiments**

1µg of total RNA was used for the reverse transcription, using the QuantiTect Reverse Transcription kit (QIAGEN, Hilden, Germany), with a mix of oligo (dT) and random primers to prime first-strand synthesis. The SYBR master mix plus (Eurogentec, Liege, Belgium) was used for qPCR. The mRNA level was normalized to 36B4 mRNA.

| **Gene Name** | **Species** | **Oligonucleotide Sequence (5' to 3')** |
|---|---|---|
| ATF4 | *Homo Sapiens* | Forward ATGACCGAAATGAGCTTCCTG<br>Reverse GCTGGAGAACCCATGAGGT |
| PHGDH | *Homo Sapiens* | Forward CTGCGGAAAGTGCTCATCAGT<br>Reverse TGGCAGAGCGAACAATAAGGC |
| PSAT1 | *Homo Sapiens* | Forward TGCCGCACTCAGTGTTGTTAG<br>Reverse GCAATTCCCGCACAAGATTCT |
| PSPH | *Homo Sapiens* | Forward GAGGACGCGGTGTCAGAAAT<br>Reverse GGTTGCTCTGCTATGAGTCTCT |
| SHMT1 | *Homo Sapiens* | Forward CTGGCACAACCCCTCAAAGA<br>Reverse AGGCAATCAGCTCCAATCCAA |
| SHMT2 | *Homo Sapiens* | Forward CCCTTCTGCAACCTCACGAC<br>Reverse TGAGCTTATAGGGCATAGACTCG |
| 36B4 | *Homo Sapiens* | Forward CAGATTGGCTACCCAACTGTT<br>Reverse GGCCAGGACTCGTTTGTACC |

**Colony Formation Assay**

The cells were seeded (500 cells per condition) in 60 diameters dishes and treated or not with sunitinib (sun, 0.5 μM) and/or NCT-503 (NCT, 30 μM). Colonies were detected after 10 days of culture. Cells were then washed, fixed and colored at room temperature for 20min with Crystal Violet. Then, the colony area was measured by ImageJ. Three different experiments were performed.

**Scratch assay / migration assay**

The cells were seeded in 12 well plates (180,000 cells per condition). 24 hours later, each well was artificially wounded by scratching the cell monolayer with a 20-μl plastic pipette tip. Images of the scratch wounds were taken 16 hours after wounding and the measured wound width was subtracted from the wound width at time zero to obtain the net wound closure. The wound closure areas were measured with ImageJ. The experiments were repeated at least three times.

**Spheroid formation and invasion assay**

For spheroid generation, 500 µl/well of cell suspensions at 10 000 cells / well were dispensed into agar-coated 24 wells plates (1.5% of agarose). After 3 days of spheroids initiation, siRNA transfection was performed (25nM of sicontrol or siPSAT1), then, one day after, spheroids were included in Matrigel (1mg/ml) and 500µl of DMEM media with 7% FCS were added on the top. For NCT-503 (NCT, 30 µM) and sunitinib (sun, 2.5 µM), treatments were performed the day of inclusion in Matrigel both in the Matrigel and in the media on the top.

To follow invasion, area invasion was measured at different days (0, 3 days and 7 days).

**Metabolite profiling for targeted steady state and flux analyses**

To determine the relative abundances of intracellular metabolites, extracts were prepared and analyzed by LC-MS/MS. Briefly, for targeted steady-state samples, metabolites were extracted on dry ice with 4mL 80% methanol (−80°C), as described previously (Weinberg et al., 2019; Yuan et al., 2019). Insoluble material was pelleted by centrifugation at 3000 g for 5 min, followed by two consecutive extractions of the insoluble pellet with 0.5-ml 80% methanol, with centrifugation at 20,000 g for 5 min. The 5-ml metabolite extract from the pooled supernatants was dried down under nitrogen gas using the N-EVAP (Organomation, Inc, Associates). 50% acetonitrile was added to the samples for reconstitution following by vortexing for 30 sec. Samples solution was centrifuged 20,000 g for 30 min at 4 °C. Supernatant was collected for LC-MS analysis. For isotope tracing experiments, cells were seeded in biological triplicate (~80% confluent), washed once with serum-free DMEM and then incubated in the glucose deprived DMEM (US Biological, D9800-02) containing 25 mM of $^{13}C_6$-glucose (Cambridge Isotope Laboratories, CLM-8367-PK) for 16 hours and metabolites were extracted. For both, steady-state and tracing experiments, samples were analyzed by High-Performance Liquid Chromatography and High-Resolution Mass Spectrometry and Tandem Mass Spectrometry (HPLC-MS/MS).. In detail, the LC-MS/MS system is comprised of a Thermo Q-Exactive in line with an electrospray ionization (ESI) source and an Ultimate3000 (Thermo) series HPLC consisting of a binary pump, degasser, and auto-sampler outfitted with a Xbridge Amide column (Waters; dimensions of 2.3 mm × 100 mm and a 3.5 µm particle size). The mobile phase A contained 95% (vol/vol) water, 5% (vol/vol) acetonitrile, 10 mM ammonium hydroxide, 10 mM ammonium acetate, pH = 9.0; B was 100% Acetonitrile. The gradient was as following: 0 min, 15% A; 2.5 min, 30% A; 7 min, 43% A; 16 min, 62% A; 16.1–18 min, 75% A; 18–25 min, 15% A with a flow rate of 150 µL/min. The capillary of the ESI source was arranged for 275 °C, with sheath gas at 35 arbitrary units, auxiliary gas at 5 arbitrary units and the spray voltage at 4.0 kV. In positive/negative polarity switching mode, an *m*/*z* scan range from 60 to 900 was chosen and MS1 data was collected at a resolution of 70,000. The automatic gain control (AGC) target was set at $1 \times 10^6$ and the maximum injection time was 200 ms. The top 5 precursor ions were then fragmented, in a data-dependent manner, using the higher energy collisional dissociation (HCD) cell set to 30% normalized collision energy in MS2 at a resolution power of 17,500. Besides matching m/z, metabolites are identified by matching either retention time with analytical standards and/or MS2 fragmentation pattern. Data acquisition and analysis were carried out by Xcalibur 4.1 software and Tracefinder 4.1 software, respectively (both from Thermo Fisher Scientific). Metabolomic Pathway Analysis was calculated using MetaboAnalyst software (https://www.metaboanalyst.ca/) (Chong et al., 2019).

**Ectopic model of RCC**

Five million 786-O cells were injected subcutaneously into the flank of 5-week-old nude (nu/nu) female mice (Janvier). The tumor volume was determined with a caliper ($v = L \times l^2 \times 0.5$). When the tumor reached 100 mm$^3$, mice were treated 5 days a week for 4 weeks, by gavage with placebo (dextrose water vehicle) or sunitinib (40 mg/kg). This study was carried out in strict accordance with the recommendations in the Guide for the Care and Use of Laboratory Animals.

**siRNA assay**

Cells were transfected with either 25 nmol/L of siPSAT1 (smartpool- Dharmacon) or si-Control (smartpool- Dharmacon) by transfection with Lipofectamine RNAiMAX (Invitrogen) in Opti-MEM.

**Enzymatic dissociation of primary**

Tissue dissection, enzymatic digestion (DNase, collagenase and dispase, 37°, 30min), and mechanical dissociation were performed to obtain single-cells culture of primary RCC cells. Cells were then grown with PromoCell Renal Epithelial Cell Growth Medium 2 (PromoCell).

**Gene expression microarray analysis**

Normalized RNA sequencing data produced by The Cancer Genome Atlas (TCGA) were downloaded from cBiopotal (www.cbioportal.org, TCGA Provisional; RNA-Seq V2). Data were available for 538 RCC tumor samples TCGA subjected to mRNA expression profiling (Firehose legacy). The Kaplan–Meier method was used to produce overall survival and progression free survival curves.

**CRISPRCas9 generation**

The selected gRNA target regions were designed using the ChopChop web tool ([2]). gRNA guides used in this study are:

CrPSAT1_ex6 GTCATCACGGACAATCACCA (#5)

CrPSAT1_ex6 GACCTTGTATTCCAGGACCGA (#17)

CrPSAT1_ex2 GACTCAGTAAGTCCCCGCGAG (#6)

Bold G in 5' is added due to the transcription initiation requirement of a 'G' base for human U6 promoter. 786-O WT cells were transfected using PEI (Polyplus transfection) with pSpCas9(BB)-2A-GFP (PX458) plasmids (a gift from Feng Zhang (plasmid #48138, Addgene, Watertown, MA, USA) ([1]) containing CRISPR-Cas9 and guide RNA.

As the pSpCas9(BB)-2A-GFP (pX458) plasmid used in the study contains GFP, single-cell sorting was conducted (24-h post transfection) using BD FACS Melody (BD Biosciences, Franklin Lakes, USA). Individual clones were cultivated in DMEM 7% FBS. Each individual clone was analyzed for PSAT1 expression by RT-QPCR and immunoblot.

**Statistical analysis**

One-way ANOVA followed by Tukey's post hoc tests were performed in GraphPad Prism 7.0 to determine differences between each group when more than two conditions were present. A two-tailed Student's t tests were performed for two pairwise comparisons. All error bars represent standard deviation (SD). A value of $p < 0.05$ was considered significant.
